# A therapeutic role for a regulatory glucose transporter1 (Glut1)-associated natural antisense transcript

**DOI:** 10.1101/2025.03.26.644647

**Authors:** Maoxue Tang, Sasa Teng, Yueqing Peng, Ashley Y. Kim, Peter Canoll, Jeffrey N. Bruce, Phyllis L. Faust, Kailash Adhikari, Darryl C. De Vivo, Umrao R. Monani

## Abstract

The mammalian brain relies primarily on glucose for its energy needs. Delivery of this nutrient to the brain is mediated by the glucose transporter-1 (Glut1) protein. Low Glut1 thwarts glucose entry into the brain, causing an energy crisis and, triggering, in one instance, the debilitating neurodevelopmental condition – Glut1 deficiency syndrome (Glut1DS). Current treatments for Glut1DS are sub-optimal, as none address the root cause – low Glut1 – of the condition. Levels of this transporter must respond rapidly to the brain’s changing energy requirements. This necessitates fine-tuning its expression. Here we describe a long-noncoding RNA (lncRNA) antisense to *Glut1* and show that it is involved in such regulation. Raising levels of the lncRNA had a concordant effect on *Glut1* in cultured human cells and transgenic mice; reducing levels elicited the opposite effect. Delivering the lncRNA to Glut1DS model mice via viral vectors induced *Glut1* expression, enhancing brain glucose levels to mitigate disease. Direct delivery of such a lncRNA to combat disease has not been reported previously and constitutes a unique therapeutic paradigm. Moreover, considering the importance of maintaining homeostatic Glut1 levels, calibrating transporter expression via the lncRNA could become broadly relevant to the myriad conditions, including Alzheimer’s disease, wherein Glut1 concentrations are perturbed.

## Introduction

Although it accounts for just 2% of the weight of the average adult, the human brain has an unusually high energy requirement, consuming ∼25% of all nutrient intake (1). This energy is supplied to the brain mainly in the form of glucose and delivered to the cerebral parenchyma via the glucose transporter1 (Glut1) protein (2). Complete loss of Glut1 causes death *in utero* in mice(3) and is almost certainly embryonic lethal in humans too. In contrast, low (∼ 50%) Glut1, while sufficient to ensure embryonic development, causes an energy crisis – neuroglycopenia – which is particularly detrimental to the brain and CNS (4, 5). Understanding and combating neuroglycopenia is broadly relevant, as it characterizes several common conditions including Alzheimer’s disease (AD) and traumatic brain encephalopathies (TBEs) (6). Yet, it is perhaps best investigated in and exemplified by the neurodevelopmental disorder, Glut1 deficiency syndrome (Glut1DS) (7). Resulting from *SLC2A1 (Glut1)* haploinsufficiency – mostly a consequence of *de novo* mutations – Glut1DS is reported to occur as frequently as 1 in 24,000 newborns(8–10). Individuals afflicted with the condition classically become symptomatic during infancy, exhibiting a complex phenotype that initially manifests as abnormal eye-head movements, intractable epileptic seizures and neurodevelopmental delay (11). A complex movement disorder that combines elements of ataxia, dystonia and spasticity often develops later in life and can be complicated by exercise-induced dyskinesia and hemolytic anemia (7).

Although the genetic defect underlying Glut1DS was identified more than two decades ago, there is still no treatment that directly addresses Glut1 deficiency. Ketogenic diets, that supply the brain with an alternate albeit imperfect energy source, ketone bodies, are the current mainstay for individuals diagnosed with the condition (7). Such diets mitigate seizure activity in patients but often fail to prevent movement disorders and, in some instances, trigger serious adverse effects, e.g., significant reduction of bone mass, long-term cardiovascular complications due to atherosclerosis and, in rare instances, the induction of a state of coma (12–14). Glut1DS thus remains a disease with an urgent unmet medical need.

Considering the link between *SLC2A1* haploinsufficiency and Glut1DS, one intuitively attractive alternative therapeutic strategy to ketogenic diets that directly addresses the molecular defect in the condition is simply to restore cerebral levels of the transporter. This may be accomplished with small molecules that relieve transcriptional or translational repression but may equally well be brought about by regulatory RNAs. This study focuses on the latter approach and stems from the discovery of a novel RNA located in the *Glut1* locus. The RNA, a natural antisense long non-coding transcript (lncRNA) has a potent effect on *Glut1* expression. Using a combination of cultured human cells, human tissues, transgenic mice and a well-established rodent model of Glut1DS, we show that the lncRNA concordantly regulates *SLC2A1,* increasing Glut1 to therapeutic levels. Indeed, delivering the transcript in a viral vector to model mice raised cerebral Glut1 sufficiently to prevent the onset of motor defects, ameliorate abnormal brain activity and suppress the neuroinflammation commonly seen in mutants. This therapeutic modality wherein a natural antisense transcript is employed to stimulate expression of its protein-coding cognate to treat disease is novel. Moreover, considering the myriad conditions involving reduced Glut1, the therapeutic strategy described here for Glut1DS could become relevant beyond just this rare neurodevelopmental disorder.

## Results

### A natural antisense transcript (NAT) concordantly regulates human SLC2A1 (Glut1) expression

While examining the human *Glut1* locus, we discovered an expressed sequence located 5’ to the *SLC2A1* gene. On closer inspection, the expressed sequence turned out to be a spliced, polyadenylated 1.1kb natural antisense lncRNA transcript consisting of 4 exons spread over ∼24kb of genomic sequence on human Chr.1 (Fig. 1A). Considering a growing recognition that such lncRNAs regulate expression of their cognate protein-coding genes(15), we inquired if the transcript – now designated *SLC2A1-DT* in public databases – modulated *Glut1* activity. To do so, we first knocked down lncRNA expression using siRNAs (Fig. S1A). One of these (siRNA#2) was subsequently employed to examine the effect of lncRNA knockdown on Glut1 levels (Fig. 1B). Natural antisense transcript (NAT) expression often exhibits a discordant relationship with that of its sense strand partner (16). We were therefore surprised to find that knockdown of the lncRNA produced a concordant effect, resulting in significantly lower Glut1 levels. Consistent with reduced *Glut1* RNA levels, Glut1 protein also decreased (Figs. 1B – D). Conversely, overexpressing lncRNA from a plasmid in human fibroblasts raised Glut1 RNA and protein (Figs. 1E, F), suggesting that the element expressing the transcript need not be positioned in *cis* to bring about an effect. These effects of the *SLC2A1-DT* lncRNA on *Glut1* were also observed in human brain endothelial cells (BECs), which express abundant transporter and are important in delivering glucose to the brain parenchyma. Indeed, knocking down and raising lncRNA expression respectively inhibited and enhanced *Glut1* transcript levels significantly (Figs. S1B – D). Finally, consistent with the ability to function in *trans*, co-transfecting the lncRNA into cultured cells with a construct containing a 6kb Glut1 promoter fragment driving luciferase stimulated expression from the reporter (Figs. S1E, F).

**Figure 1.**
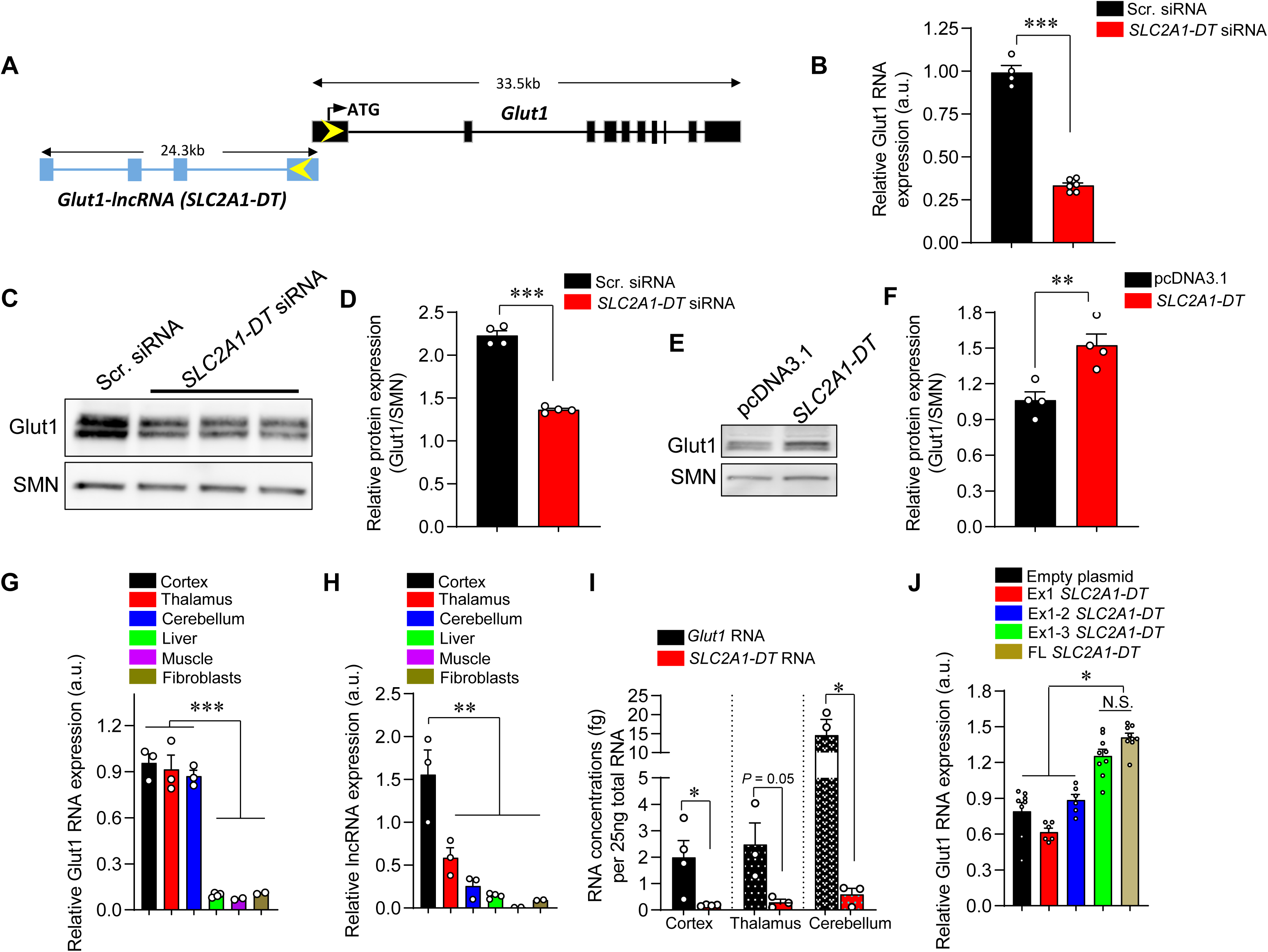
A natural antisense transcript regulates *SLC2A1 (Glut1)* expression concordantly. **(A)** Schematic depicting spatial relationship between the *Glut1* gene and the *SLC2A1-DT* natural antisense transcript. Exons 1 of *Glut1* and *SLC2A1-DT* overlap over 128bp. **(B)** Quantified results of *SLC2A1-DT* knockdown on Glut1 RNA levels in human fibroblasts. ***, *P* < 0.001, *t* test, n ≥ 4 replicates for each condition. **(C)** Representative western blot showing effect of *SLC2A1-DT* knockdown on Glut1 protein levels in fibroblasts. **(D)** Quantified results of western blots to determine effect of suppressing *SLC2A1-DT* expression on Glut1 protein levels. ***, *P* < 0.001, *t* test, n = 4 replicates for each condition. **(E)** Representative western blot depicting increase in Glut1 protein in human fibroblasts following over-expression of the *SLC2A1-DT* lncRNA. **(F)** Graph depicting quantified increase of Glut1 protein when the lncRNA is over-expressed. **, *P* < 0.01, *t* test, n = 4 replicates each. Expression profiles of **(G)** Glut1 and **(H)** lncRNA in various human tissues. **, ***, *P* < 0.01 and *P* < 0.001 respectively one-way ANOVA. **(I)** Graph depicting low levels of the lncRNA relative to Glut1 levels in human brain tissues sampled. *, *P* < 0.05, *t* tests, n ≥ 3 samples. **(J)** Quantified relative Glut1 RNA levels obtained from experiments to determine the minimum *SLC2A1-DT* exons to promote *Glut1* activity. Note: *, *P* < 0.05, Kruskal-Wallis test, n ≥ 6 test replicates with each construct.

NATs are typically expressed at low levels, with a tissue specificity reflective of that of their sense strand partners (16, 17). Accordingly, to further characterize *SLC2A1-DT*, we examined its expression and that of *Glut1* in diverse human tissue samples. Akin to high Glut1 levels in brain samples, the lncRNA was expressed especially robustly in cerebral cortex (Figs. 1G, H). This pattern of concordant expression was also observed in two human brain tumor samples wherein characteristically and abnormally high *Glut1* transcripts were accompanied by enhanced lncRNA expression – relative to levels in healthy brain tissue (Figs. S1G, H). Still, consistent with generally low expression of lncRNAs, *SLC2A1-DT* was found at markedly lower concentrations relative to Glut1 in the human brain regions that we examined (Fig. 1I). In a final set of studies using cultured cells, we inquired if truncated versions of the lncRNA retained the ability to regulate *Glut1.* In parallel, we investigated the cellular localization of the transcript. We found that exons 1-3 of the lncRNA were just as effective in raising Glut1 as the full-length transcript was, whereas constructs containing just exon 1 or the first two exons failed to stimulate *Glut1* expression (Fig. 1J). Experiments to determine the cellular distribution of the lncRNA demonstrated that it localizes to the cytoplasm as well as nucleus (Fig. S1I). Collectively, these results suggest that *SLC2A1-DT*, a recently annotated NAT in the human *SLC2A1* locus, not only regulates *Glut1* expression in a concordant manner but can also do so in *trans*.

### Transgenic expression of SLC2A1-DT mitigates disease in Glut1DS model mice

Considering the stimulatory effect of the *SLC2A1*-*DT* NAT on *Glut1* expression and our long-term objective of developing therapies for Glut1DS that restore transporter levels to patients afflicted with the condition, we inquired if the NAT also induced *Glut1* expression in the whole organism. To do so, we generated mice which were transgenic for a genomic fragment harboring the human *Glut1* locus. Serendipitously, one of the resulting lines was found to contain a truncated fragment harboring only part of the *SLC2A1* gene but the entirety of the *SLC2A1-DT* NAT (Fig. S2A). PCR analysis of material from brain tissue isolated from mice from this line demonstrated that the NAT was indeed expressed in the line (Fig. 2A and Fig. S2B). Interestingly, this resulted in an increase in murine *Slc2a1* (Fig. 2B and Fig. S2C), suggesting modulatory function of the NAT across species. To determine if the NAT-mediated increase in murine *Glut1* was of physiological consequence, we introduced the transgene bearing *SLC2A1-DT* into a well-established mouse model of Glut1DS, which is haploinsufficient for *Slc2a1 (Glut1^+/-^)* (3). Glut1 deficient mice harboring the transgene *(Glut1^+/-^;SLC2A1-DT^tg^)* and relevant controls were then assessed three different ways. In a test of motor activity on a rotarod, we found that *Glut1^+/-^;SLC2A1-DT^tg^* mutants performed significantly better than mutants absent the lncRNA but less well than wild-type (WT) controls *(Glut1^+/+^)* (Fig. 2C). Consistent with the improved motor performance, we found that hypoglycorrhachia [low cerebrospinal fluid (CSF) glucose], a signature feature of Glut1DS, was mitigated in *Glut1^+/-^;SLC2A1-DT^tg^*mutants (Fig. 2D). As blood glucose levels in these mutants were unchanged by expression of the transgene, CSF:blood glucose ratios were also significantly enhanced (Fig. 2E). Hypolactorrhachia (low CSF lactate), an additional characteristic of Glut1DS (7) was also ameliorated in *Glut1^+/-^;SLC2A1-DT^tg^* mutants, with lactate levels in the mice, in these experiments, raised to those observed in WT controls (Fig. S2C). Glut1DS is characterized by debilitating seizures. In model mice, this phenotype manifests as abnormal spike-wave discharges (SWDs) observed in electro-encephalograms (EEGs) (7, 18). We found that the number of SWDs measured over 24hrs in mutants expressing *SLC2A1-DT* was reduced relative to cohorts devoid of the NAT and no different from those in WT controls (Fig. 2F). In aggregate, these findings suggest 1) that the ability of the *SLC2A1-DT* lncRNA to modulate *Glut1* expression in cultured cells is maintained in the intact organism – a rodent in this instance, 2) that it is indeed able to function in *trans*, as the transgene expressing the lncRNA was mapped to an altogether different region of the mouse genome (Chr.12qC2) from mouse *Slc2a1* (Chr.4qD2.1) and, finally 3) that the ability of the lncRNA to raise *Glut1* expression is of physiological consequence, reflected in amelioration of the Glut1DS phenotype of model mice.

**Figure 2.**
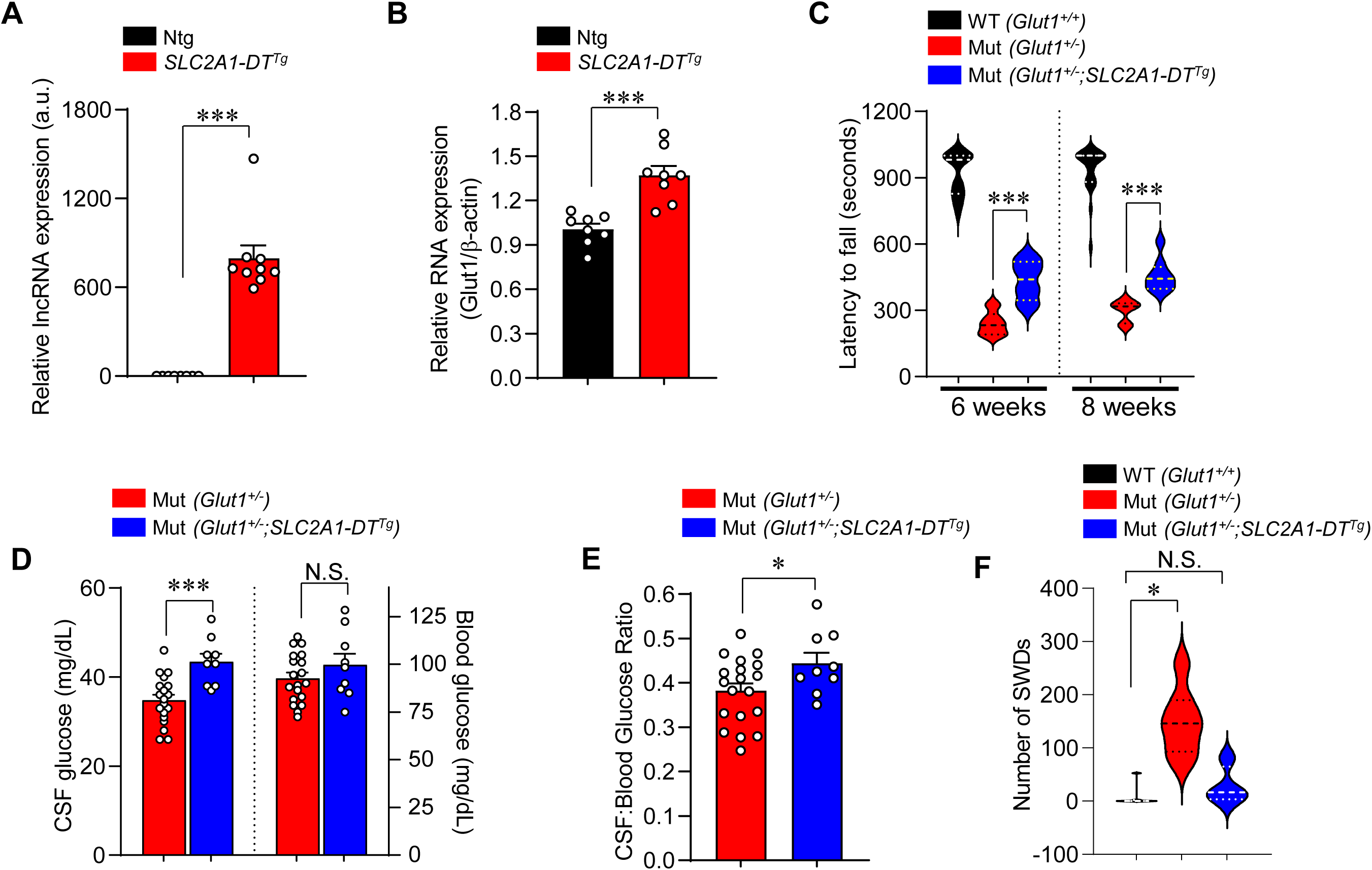
Transgenic expression of the *SLC2A1-DT* lncRNA suppresses Glut1DS in model mice. Quantified expression of **(A)** the lncRNA and **(B)** murine *Glut1* in transgenic mice and non-transgenic littermates. ***, *P* < 0.001, *t* test, n ≥ 8 mice of each cohort. **(C)** Graph depicting significantly improved motor performance of Glut1DS mutants harboring the lncRNA transgene. ***, *P* < 0.001, *t* test, n = 8 – 22 mice of each cohort. Quantified results of **(D)** CSF and blood glucose level measurements and **(E)** CSF:blood glucose ratios in mutants with or without the lncRNA transgene. *, ***, *P* < 0.05 and *P* < 0.001 respectively, *t* tests, n = 9 – 19 mice of each cohort. **(F)** Graph depicts fewer seizures manifesting as spike-wave discharges in Glut1DS mutants bearing the lncRNA transgene relative to mutants devoid of the transgene. *, *P* < 0.05, Mann-Whitney test, n = 4 – 7 mice of each cohort.

### SLC2A1-DT from an exogenous source mitigates overt disease in Glut1DS model mice

Considering its effects when expressed as a transgene and given our quest to develop novel Glut1DS therapies that address the root cause – low Glut1 – of the condition, we inquired if *SLC2A1-DT* from an exogenous source might also mitigate disease in model mice. To test this possibility, we packaged the lncRNA into a self-complementing adeno-associated viral vector (scAAV.PHP.eB) (19) and delivered it systemically at two different concentrations, 4.2 x 10^11^VGs (low dose) and 8.4 x 10^11^VGs (high dose), to postnatal 1 (PND1) Glut1DS mutants. We began by assessing the effects of such delivery on motor performance of 5-week-old lncRNA-treated mutants. AAV-eGFP-treated mutants and WT *Glut1^+/+^*mice served as controls. As expected, AAV-eGFP-treated mutants performed very poorly relative to WT controls on the rotarod. In contrast, the performance of mutants delivered either a low or high dose of the AAV-lncRNA was indistinguishable from that of *Glut1^+/+^* mice (Fig. 3A). Notably, this outcome was sustained over a 4-week period despite a subtle overall decrease in latency to fall off the rotarod – in all mouse cohorts. Differences between the mutants treated with the low or high doses of the lncRNA were mostly undetectable. We next assessed the effect of the lncRNA on hypoglycorrhachia. Measurements of CSF glucose revealed that while levels in lncRNA-treated mutants were lower than those in WT mice, they were nevertheless significantly higher than those of AAV-eGFP-treated mutants (Fig. 3B). Akin to observations in our transgenic lines, blood glucose levels remained unaltered whereas CSF:blood glucose ratios rose in lncRNA-treated versus eGFP-treated mutants (Figs. 3B, C). Glut1DS patients exhibit decelerating head growth (7). In model mice, this manifests as micrencephaly (3). Expectedly, brain sizes of AAV-eGFP-treated *Glut^+/-^* mutants were markedly lower than those of *Glut1^+/+^* controls, notwithstanding equivalent body weights. In contrast, and consonant with previous outcomes, the micrencephalic phenotype was significantly less severe in mutants treated with AAV-lncRNA (Figs. 3D, E). Differences between the cohorts treated with low or high lncRNA doses were once again undiscernible. These results suggest that *SLC2A1-DT* from an exogenous source is indeed salutary to Glut1DS mutants. Moreover, our inability to detect differences between mutants administered high or low doses of AAV-lncRNA suggested that the NAT has a ceiling effect. This ceiling appears to be reached, in the rodent model, with the low dose of the therapeutic vector.

**Figure 3.**
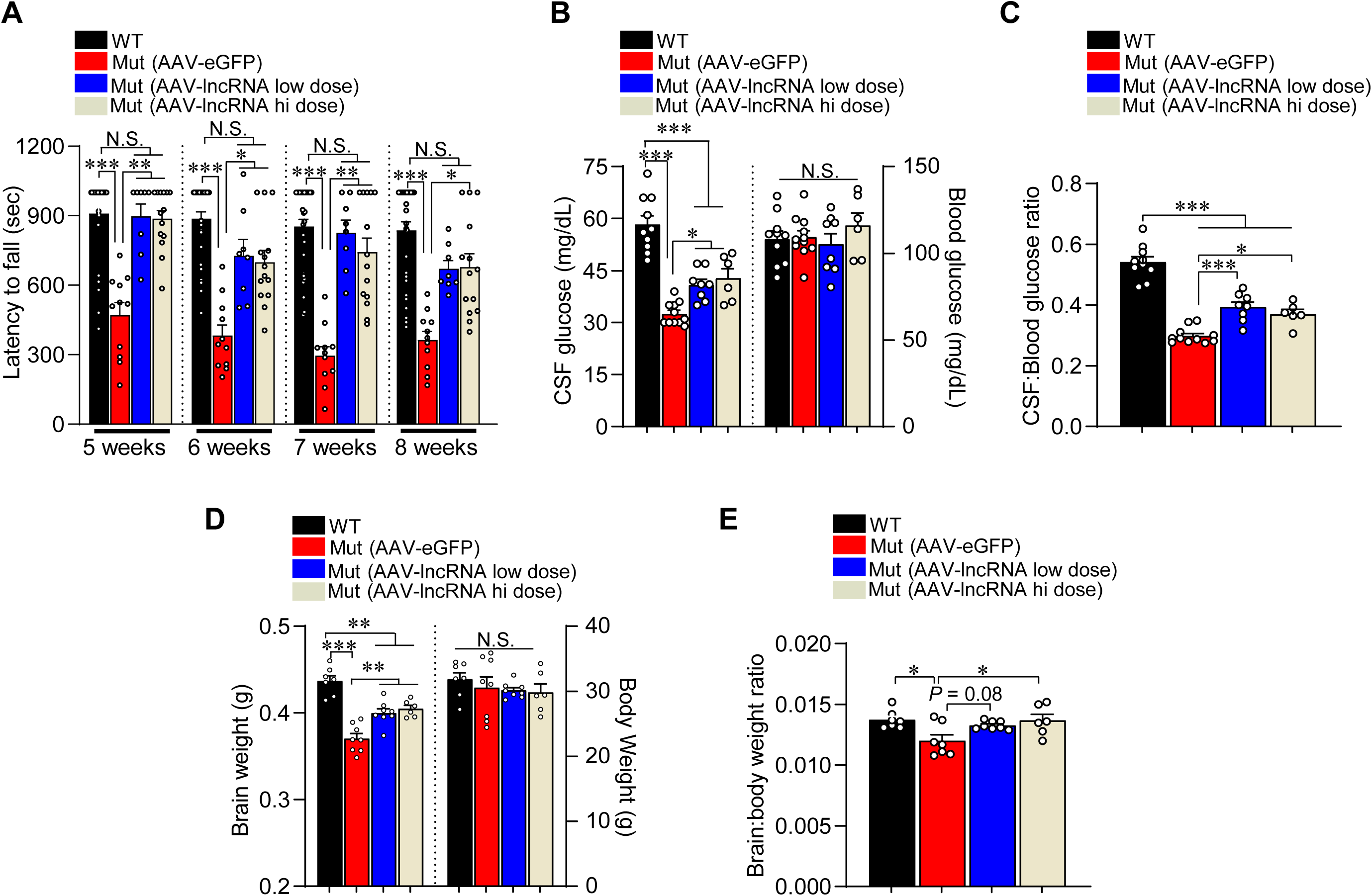
Viral vector-mediated delivery of the *SLC2A1-DT* lncRNA lessens Glut1DS disease severity in model mice. **(A)** Quantified results of rotarod motor performance tests of the indicated mouse cohorts over a 1-month time period. Note significant improvement in the performance of mutants administered the therapeutic lncRNA. *, **, ***, *P* < 0.05, *P* < 0.01 and *P* < 0.001 respectively, Kruskal-Wallis test, n = 8 – 32 mice of each cohort. Graphs depicting **(B)** CSF and blood glucose levels and, **(C)** CSF:blood glucose ratios in the various cohorts of mice. *, ***, *P* < 0.05 and *P* < 0.001 respectively, one-way ANOVA, n = 6 – 11 mice of each cohort. **(D, E)** Graphs showing evidence of reduced micrencephaly in Glut1DS mutant mice administered either low or high dose of AAV-lncRNA. *, **, ***, *P* < 0.05, *P* < 0.01 and *P* < 0.001 respectively, one-way ANOVA, n = 6 – 8 mice of each cohort. Outcomes quantified in panels B – E were assessed in 5-month-old mice.

### AAV-mediated delivery of SLC2A1-DT raises Glut1 and ameliorates brain pathology in Glut1DS mutants

To investigate the molecular basis of disease rescue in mutants administered the lncRNA, we examined levels of the transcript in the brain microvasculature fractions of these mice and controls administered the AAV-eGFP construct. Q-PCR analysis revealed robust expression of the lncRNA in mutants administered the low dose of the vector and even greater transcript levels in mutants injected with the high dose of the vector (Fig. 4A). Expectedly, AAV-eGFP-injected animals expressed negligible amounts of the human lncRNA. Next, we assessed the effect of the lncRNA on *Glut1* RNA in the various cohorts of mice. Predictably, AAV-eGFP-injected mutants expressed significantly lower Glut1 in brain capillary and neuropil fractions relative to amounts of the transcript in these compartments of WT controls (Figs. 4B). In contrast, but consistent with lncRNA measurements, *Glut1* transcript was raised in the two sets of mutants injected with AAV-lncRNA. Interestingly, Glut1 in brain capillaries but not neuropil of these mutants was completely normalized suggesting a more potent effect of the lncRNA in tissue known to express abundant Glut1 (20). Reflective of a ceiling effect of the lncRNA, Glut1 levels were equivalent in mutants administered low or high dose of the vector – in each of the brain fractions examined. Western blots to assess Glut1 protein in brain tissue of the various cohorts confirmed the RNA expression analysis; the lncRNA restored levels of the transporter to mutants administered the different doses of therapeutic vector (Figs. 4C – F).

**Figure 4.**
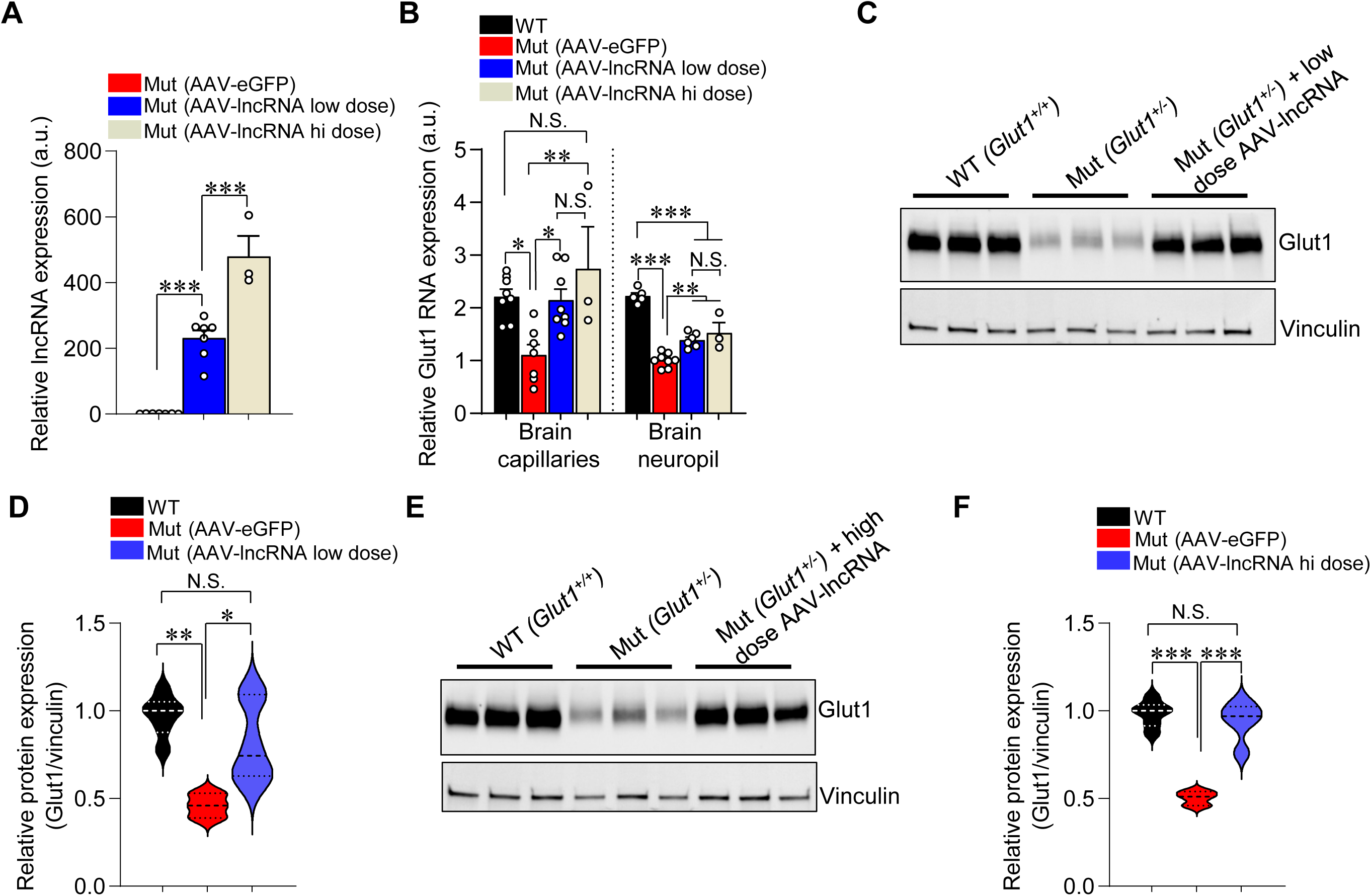
AAV-mediated delivery of the *SLC2A1-DT* lncRNA stimulates *Glut1* expression in Glut1DS model mice. **(A)** Results of Q-PCR on brain tissue of mice showing robust expression of the lncRNA in Glut1DS mutants administered either low or high dose of AAV-lncRNA. ***, *P* < 0.001, one-way ANOVA, n = 3 – 7 mice of each cohort **(B)** AAV-mediated expression of the lncRNA raises murine *Glut1* RNA levels in brain fractions of the two relevant cohorts of mutants in comparison to mutants administered an AAV-eGFP construct. *, **, ***, *P* < 0.05, *P* < 0.01 and *P* < 0.001 respectively, one-way ANOVA, n = 3 – 8 mice of each cohort. **(C)** Representative western blot and **(D)** quantified representation of murine Glut1 protein in brain blood vessels of mutants administered the *low* dose of AAV-lncRNA and relevant controls. *, **, *P* < 0.05 and *P* < 0.01 respectively, one-way ANOVA, n = 3 – 6 mice of each cohort. **(E)** Representative western blot and **(F)** graphical plot of murine Glut1 protein in cerebral blood vessels of mutants administered the *high* dose of AAV-lncRNA and relevant controls. ***, *P* < 0.001, one-way ANOVA, n = 3 – 6 mice of each cohort.

We previously showed that Glut1 haploinsufficiency in model mice triggers profound and early neuroinflammation (21). This is accompanied by arrested development of the brain microvasculature (18, 21). Accordingly, we examined brain sections of our mice immunohistochemically to investigate the effects of the lncRNA on these pathological aspects of the disease. Consistent with our prior studies, thalamic sections from mutants treated with the control vector continued to exhibit a clearly discernible neuroinflammatory response characterized by reactive astrocytes and activated microglia (Fig. 5A, B & Fig. S3). These hypertrophied glial cells were mostly absent in WT controls, and significantly reduced in numbers in mutants administered the therapeutic vector. Similar brain sections stained with labeled lectin to highlight brain capillaries revealed a diminutive brain microvasculature in mutants treated with AAV-eGFP. In comparison, mutants treated with the therapeutic vector had significantly greater densities of capillaries (Fig. 6A, B & Fig. S4A), although the extent of the brain microvasculature did not reach the WT state. In a final assessment of the effect of the lncRNA on brain pathology, we carried out EEGs on mutants administered the low dose of therapeutic vector and compared the incidence of SWDs with those in the two control cohorts. Consistent with observations made in experiments carried out on our transgenic mice, we found that seizures in AAV-lncRNA-treated *Glut1^+/-^* mutants were reduced (Fig. 6C, D). Moreover, whereas mice treated with the AAV-eGFP construct not only exhibited significantly greater numbers of SWDs but also the occasional convulsive seizure (Fig. S4B), this latter type of seizure was completely suppressed in mutants injected with the lncRNA-containing construct. These findings are consistent with other outcomes examined in mutants treated with AAV-lncRNA and constitute compelling evidence of the therapeutic effects of delivering this modulatory RNA in a viral vector to a model of Glut1DS.

**Figure 5.**
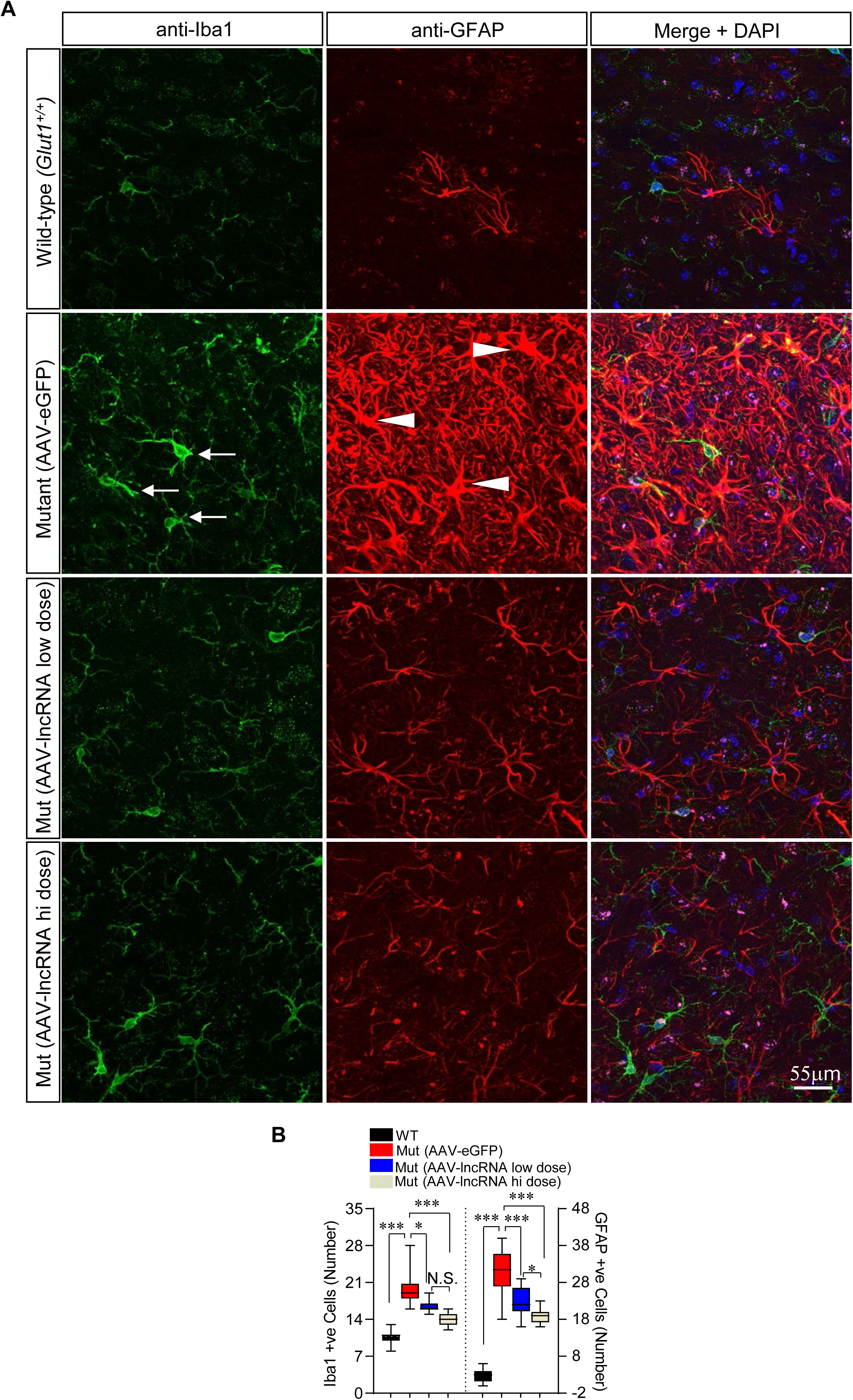
AAV-mediated delivery of the *SLC2A1-DT* lncRNA reduces neuroinflammation in Glut1DS model mice. **(A)** Representative thalamic sections from 4 – 5-month-old controls and mutants treated with the lncRNA. Pronounced neuroinflammation in AAV-eGFP-treated mutants characterized by activated, hypertrophic microglia (arrows) and reactive astrocytes (arrowheads) is observed. Fewer such cells were observed following treatment with AAV-lncRNA. **(B)** Graph depicts quantified numbers of reactive astrocytes and activated microglia in the various cohorts of mice. *, ***, *P* < 0.05 and *P* < 0.001 respectively, Kruskal-Wallis test (microglia) and one-way ANOVA (astrocytes), n = 3 non-adjacent fields from N = 3 – 6 mice of each cohort.

**Figure 6.**
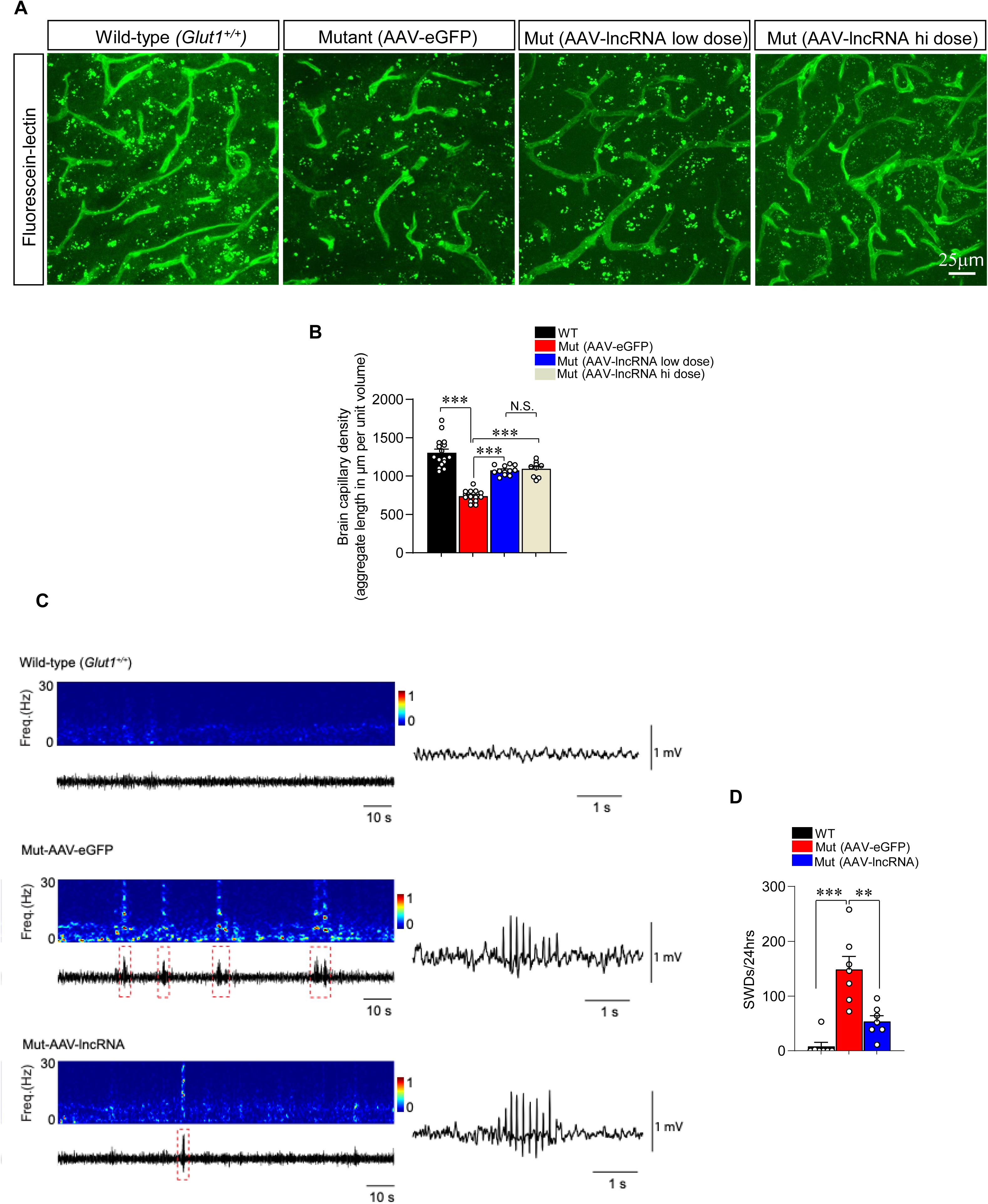
AAV-mediated delivery of the *SLC2A1-DT* lncRNA stimulates brain capillary formation and reduces seizures in Glut1DS model mice. **(A)** Representative thalamic sections from 4 – 5-month-old controls and mutants treated with the lncRNA. Sections were stained with labeled lectin to reveal brain capillaries. Note fewer capillaries in mutant treated with AAV-eGFP and a relative normalization of the microvasculature in mutants treated with AAV-lncRNA. **(B)** Graph quantifies aggregate cerebral capillary length in the mice. \*\*\**P* < 0.001, one-way ANOVA, n = 9 regions from each of N = 3 - 5 mice of each genotype examined. **(C)** Representative EEG spectrograms at 0-30Hz (colored panels) and corresponding traces (below the colored panels) depicting individual SWDs, highlighted in red boxed areas, captured over 2min or 5s – to emphasize seizure activity – in the various cohorts of mice. **(D)** Quantification of SWDs in the three cohorts of mice. **, ***, *P* < 0.01 and *P* < 0.001 respectively, Mann-Whitney tests, n = 7 mice in each group.

### Sustained therapeutic effects of AAV-mediated delivery of SLC2A1-DT to Glut1DS model mice

Durability of benefit that is accrued following gene delivery in a viral vector is critically important for its future use in the clinic. Accordingly, a subset of the various cohorts of mice generated for our experiments and treated with therapeutic or control vector were allowed to age. Twelve months following treatment, these mice were examined for evidence of sustained benefit from expression of the lncRNA. We began by examining Glut1 levels in the mice. As expected, AAV-eGFP-treated mutants continued to express low Glut1 in whole brain tissue relative to WT controls. In contrast, the comparatively higher levels of Glut1 we’d observed in mutants delivered either low or high doses of the lncRNA persisted in the 12-month-old cohorts. These animals continued to express ∼30% more Glut1 transcript than did mutants administered the control AAV (Fig. 7A) and is a likely consequence of sustained lncRNA activity from episomally maintained transcript delivered to cells by the AAV. We next assessed CSF glucose levels in the different mice. Consistent with higher brain Glut1 levels, we found that mutants expressing the lncRNA continued to exhibit greater concentrations of CSF glucose than those in mutants administered the AAV-eGFP construct; blood glucose levels remained unaltered (Fig. 7B). The combination of higher CSF glucose levels and unchanged blood glucose concentrations resulted in enhanced CSF:blood glucose ratios in the AAV-lncRNA versus AAV-eGFP-treated mutants (Fig. 7C). In a final set of assessments to determine persistence of therapeutic effect of *SLC2A1-DT*, we examined brain size in the treated mice. The micrencephalic phenotype in mutants treated with AAV-lncRNA remained less severe than it was in AAV-eGFP-treated mutants (Fig. 7D. Body weights in the various cohorts of mice were statistically indistinguishable but, consonant with measurements of brain size, brain:body weight ratios were higher in AAV-lncRNA versus AAV-eGFP-treated mutants (Fig. 7D, E). Together, these results attest to the durability of the effect of the lncRNA delivered to Glut1DS model mice in a viral vector.

**Figure 7.**
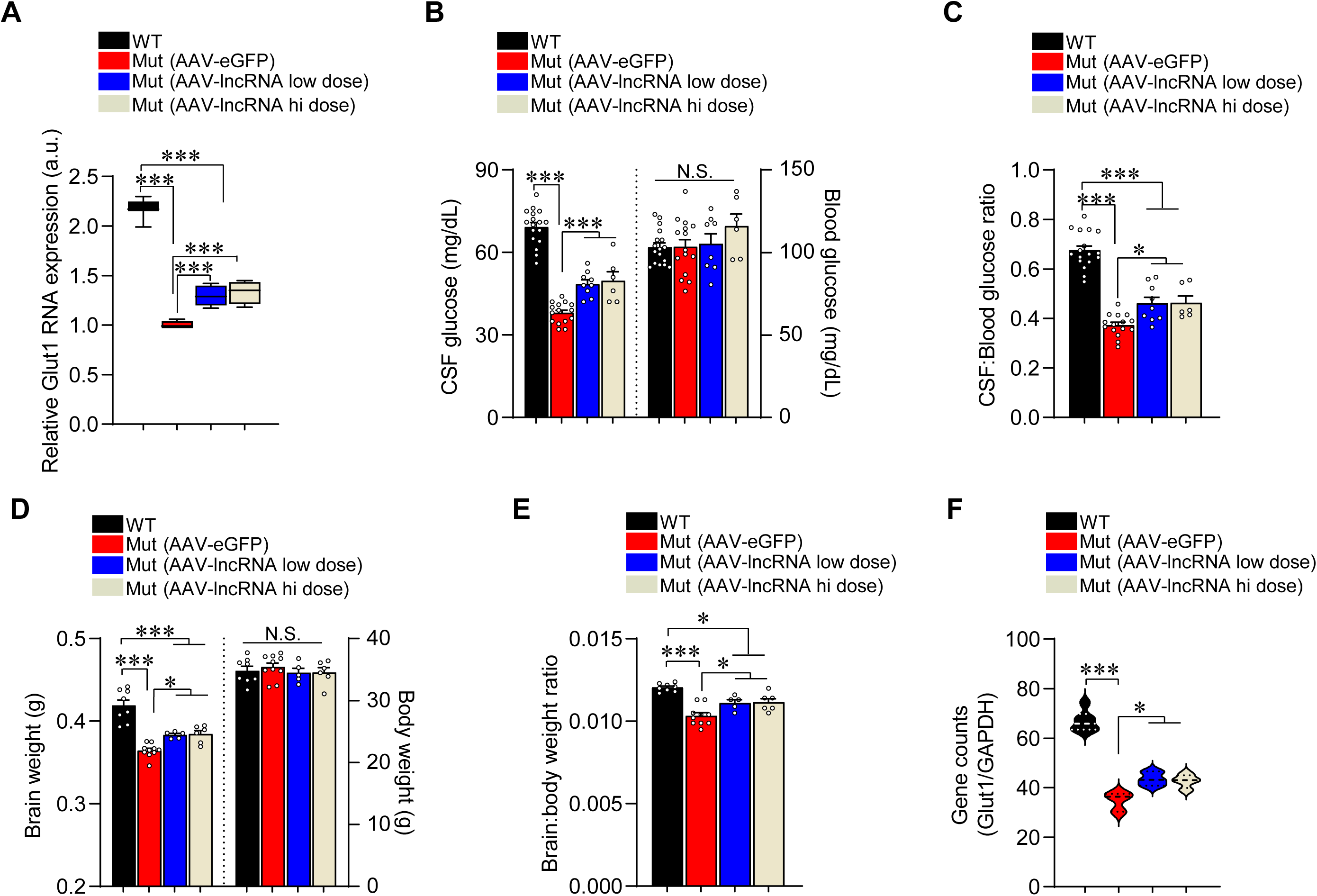
Sustained therapeutic effects deriving from AAV-mediated delivery of the *SLC2A1-DT* lncRNA. **(A)** Glut1 expression in brain tissue of Glut1DS mutants treated with the AAV-lncRNA vector remains significantly higher than it is in mutants treated with the AAV-eGFP construct. ***, *P* < 0.001, one-way ANOVA, n = 4 – 9 mice in each group. **(B)** Quantified levels of CSF and blood glucose levels depict significantly higher levels of the former in mutants treated with the AAV-lncRNA vector relative to concentrations in mutants treated with the AAV-eGFP control vector. Blood glucose levels were equivalent in all four mouse cohorts. ***, *P* < 0.001, one-way ANOVA, n = 6 – 17 mice in each group. **(C)** Quantified CSF:blood glucose ratios reflect higher levels of CSF glucose levels in AAV-lncRNA-treated mutants vis-à-vis AAV-eGFP-treated model mice. *, ***, *P* < 0.05 and *P* < 0.001 respectively, one-way ANOVA, n = 6 – 17 mice of each cohort. The micrencephalic phenotype remains less severe in Glut1DS mutants expressing the lncRNA compared to the condition in mutants injected with the AAV-eGFP vector as assessed by measuring **(D)** brain and body sizes and **(E)** calculating brain:body weight ratios. *, ***, *P* < 0.05 and *P* < 0.001 respectively, one-way ANOVA, n = 5 – 10 mice of each cohort for the two analyses. **(F)** Gene counts from RNA-Seq analysis of the various cohorts of mice illustrates sustained higher expression of Glut1 in mutants treated with AAV-lncRNA compared to mutants treated with the control AAV. *, ***, *P* < 0.05 and *P* < 0.001 respectively, one-way ANOVA, n = 3 – 6 mice of each cohort.

In anticipation of developing the lncRNA into a clinical therapeutic for Glut1DS or other conditions characterized by low Glut1 protein, and to gain insight into its mode of action, we surveilled the transcriptomes of brain tissue from 12-month-old Glut1DS mutants administered AAV-lncRNA as pups. The resulting RNA-Seq data was then analyzed for toxicological processes using the Tox Analysis function of Qiagen’s Ingenuity Pathway Analysis (IPA) tool. This indicated that the expression of the lncRNA is not significantly associated with any toxicity. Moreover, RNA-Seq analysis of WT mice and mutants administered either dose of the lncRNA revealed a total of 44 significantly differentially expressed transcripts between the mutants and controls (Table S1). A similar comparison of the transcriptomes of WT mice and mutants treated with AAV-eGFP uncovered 56 differentially expressed genes (DEGs) (Table S2). Interestingly, except for *Slc2a1 (Glut1)* none of the genes that were perturbed in this second analysis were present in the list of DEGs compiled from the comparison of WT animals and lncRNA-treated mutants (Tables S1 & S2), suggesting that at least a subset of genes that were dysregulated in *Glut1^+/-^*mutants owing to low Glut1 were normalized in expression following treatment with the lncRNA. As a specific example, we highlight TXNIP, a known interactor and facilitator of Glut1 activity (22), which was significantly altered in AAV-eGFP-treated mutants but normalized in expression by the lncRNA (Gene counts: WT=26.87±2.54; Mut-AAV-eGFP=18.34±0.20; Mut-AAV-lncRNA=25.57±1.34, *P* < 0.05, Mut-AAV-eGFP vs. WT and Mut-AAV-eGFP vs. Mut-AAV-lncRNA, one-way ANOVA, n = 3 – 6 mice of each cohort). Moreover, consonant with Q-PCR of *Glut1* expression in the various cohorts, the difference in *Glut1* transcripts between WT mice and mutants administered the therapeutic vector was smaller than it was when WT mice were compared to AAV-eGFP-treated mutants (Fig. 7A, F and Tables S1, S2). Notably, several DEGs that were revealed by comparing WT mice and AAV-eGFP-treated mutants were also present in the list of 198 DEGs catalogued by comparing mutants treated with either lncRNA or the eGFP construct (Table S3), suggesting that dysregulation of at least some of these are a result of eGFP over-expression rather than low Glut1. Gene ontology (GO) analysis of DEGs in WT versus AAV-eGFP-treated mutants to identify biological processes perturbed in Glut1DS revealed several involved in maintaining neuronal health (Table 1). Interestingly, but consistent with the Glut1-inducing effect of *SLC2A1-DT*, similar GO analysis of DEGs in WT versus lncRNA-treated mutants failed to reveal perturbations of any of these processes, suggesting once again that the lncRNA normalizes processes deranged by low Glut1. To complement the IPA analysis and ensure that long-term expression of the *SLC2A1-DT* lncRNA is benign, we carried out a histological study of the major organ systems of 12-month-old mutants treated as pups with AAV-lncRNA. A comparison of hematoxylin-eosin-stained sections of the various organs from these mutants with those of controls failed to reveal any gross cellular or morphological abnormalities associated with the sustained expression of the lncRNA (Fig. S5). These results support the IPA analysis and once again suggest that persistent expression of the lncRNA is safe. This raises optimism for future applications of the transcript for clinical therapies involving neuroglycopenia.

**Table 1.** Differentially expressed genes in Glut1DS mutants.

| GO Term | Description | P-value | Enrichment (N, B, n, b) |
| --- | --- | --- | --- |
| GO:0033693 | Neurofilament bundle assembly | 2.15E-7 | 165.43 (20183, 3, 122, 3) |
| GO:0045110 | Intermediate filament bundle assembly | 7.41E-6 | 70.90 (20183, 7, 122, 3) |
| GO:0048522 | Positive regulation of cellular process | 1.18E-5 | 1.69 (20183, 5289, 122, 54) |
| GO:0042552 | Myelination | 1.28E-5 | 11.68 (20183, 85, 122, 6) |
| GO:0008366 | Axon ensheathment | 1.46E-5 | 11.41 (20183, 87, 122, 6) |
| GO:0007272 | Ensheathment of neurons | 1.46E-5 | 11.41 (20183, 87, 122, 6) |
| GO:0060052 | Neurofilament cytoskeleton organization | 2.51E-5 | 49.63 (20183, 10, 122, 3) |
| GO:0061564 | Axon development | 2.68E-5 | 22.82 (20183, 29, 122, 4) |
| GO:0045105 | Intermediate filament polymerization/depolymerization | 3.62E-5 | 165.43 (20183, 2, 122, 2) |
| GO:0048518 | Positive regulation of biological process | 4.42E-5 | 1.58 (20183, 5963, 122, 57) |
| GO:0048523 | Negative regulation of cellular process | 6.59E-5 | 1.69 (20183, 4602, 122, 47) |
| GO:0032536 | Regulation of cell projection size | 7.47E-5 | 35.45 (20183, 14, 122, 3) |

**Note:** Analysis conducted on DEGs with $P_{adj}$ values < 0.1. For enrichment, N=total number of assessed genes examined in the RNA-Seq, B=number of genes associated with a specific GO term, n=DEGs input, b=number of genes in intersection. Enrichment=(b/n) ÷ (B/N).

## Discussion

Glucose is the mammalian brain’s preferred energy source and is delivered to the neuropil via Glut1, the principal cerebral glucose transporter. Given the brain’s limited capacity to store glucose, nutrient supply to the organ must be precisely adjusted and the amounts transported into the parenchyma continuously altered to satisfy the brain’s immediate energy requirements. Calibrating Glut1 activity to respond to these needs is therefore crucial. Here, we report on a little-described natural antisense *Glut1* lncRNA transcript engaged in such regulation and examine the therapeutic potential of the transcript for conditions involving perturbed Glut1 levels. Five principal findings emerge from our study. First and foremost, we show that the lncRNA is unlikely to be a consequence of pervasive genomic transcription, instead having evolved to play a vital role in regulating its sense-strand partner, *Glut1*. Unexpectedly, the lncRNA regulates *Glut1* in a concordant fashion. Second, and equally notable, are results demonstrating that notwithstanding its location in immediate proximity to *Glut1*, indeed overlapping part of the protein-coding gene, the lncRNA can produce its effects in *trans*. A third and related intriguing observation is that the lncRNA regulates *Glut1* across species. In this instance, the human lncRNA stimulated expression of not only the human *Glut1* gene but also its murine homolog, attesting to the significance and conserved relationship between the antisense transcript and its protein-coding cognate. A fourth noteworthy finding is that the lncRNA truly appears to tune Glut1 expression; our evidence suggests that *Glut1* stimulation by the lncRNA plateaus and is subject to a ceiling effect. This precludes potential adverse outcomes resulting from over-expressing the transporter. Finally, and perhaps most significantly, we show that the lncRNA is of therapeutic value. Using an established mouse model of Glut1DS, we demonstrate that the lncRNA can induce endogenous *SLC2A1* expression and raise Glut1 levels sufficiently to mitigate disease. Our results raise the prospect of employing the lncRNA for myriad other conditions including cancer, diabetes and AD in which Glut1 levels are disrupted.

Glucose homeostasis and therefore Glut1 activity are critical to the health and viability of the organism (6). This is especially true for the mammalian brain. Consequently, it is unsurprising that *Glut1* expression is subject to intricate control. The role of non-coding transcripts in such control has become evident only relatively recently but is consistent with an emerging appreciation of the overall physiological significance of regulatory RNAs. Reports of lncRNA-mediated control of Glut1 do exist (23–32). However, most of the associated lncRNAs referenced lie outside the *Glut1* locus, and a preponderance was reported to function by antagonizing miRNAs that interact with and modulate *SLC2A1* expression. The specific effects of the NAT we describe here have never been reported. Moreover, in the only study in which *SLC2A1-DT* was described to regulate *Glut1* expression, the relationship between sense and antisense genes was, surprisingly, found to be discordant; the lncRNA suppressed *Glut1* expression (33). One explanation for the contrast in findings to those reported here is that the previous investigation focused exclusively on hepatic cell lines – tumorigenic and non-tumorigenic – whereas our findings derive from results obtained from multiple mouse and human tissues.

The concordance in expression between *SLC2A1-DT* and *Glut1* we report is unexpected, as most NATs described antagonize their sense-strand partners (16). Yet, there are numerous instances of such transcripts stimulating expression of their protein-coding cognates. Noteworthy examples in the context of CNS conditions are *BACE1-AS*, which enhances expression of the β-secretase 1 (*BACE1*) gene implicated in AD (34), *RAB11b-AS1*, which concordantly regulates *RAB11b*, a vesicular trafficking protein linked to autism spectrum disorder (35), and *UCHL1-AS1*, which stimulates *UCHL1* expression, a ubiquitin-hydrolase-encoding Parkinson’s disease susceptibility gene (36). Other established NATs, identified in the context of cancer, that induce their sense-strand partners include *Khsp1* and *ZNF667-AS1* (37, 38). *SLC2A1-DT* and *Glut1* add to this list. Importantly, considering a growing interest in the therapeutic value of non-coding RNAs and our quest to address the root cause of Glut1DS, our results also serve as the basis of developing *SLC2A1-DT* clinically. The physiological significance of this lncRNA transcript stems from several observations including its ability to elicit an effect on *Glut1* across species and in *trans*. Consistent with these effects, we note that similarly positioned antisense transcripts have been identified in all mammals surveyed thus far (Table S4). Interestingly, birds, as represented by the domestic chicken, are devoid of such a transcript, suggesting that the lncRNA emerged relatively recently in evolution. This is consistent with the larger, more sophisticated mammalian brain and thus the need for a more refined means of catering to its energy requirements.

Despite a growing interest in the therapeutic value of regulatory RNAs (39), we are unaware of RNA therapies that employ lncRNAs. Proof-of-concept studies to stimulate gene expression by NAT-type molecules have been carried out but these have involved lncRNA mimics rather than the unaltered transcripts (40, 41). In this respect, the work we describe here is substantially novel. In addition to directly addressing the underlying cause – low Glut1 – of Glut1DS and thus providing an alternative to ketogenic diets for the condition, the therapeutic modality described here has significant advantages over *Glut1* gene replacement, which is prone to trigger Glut1 over-expression. Such over-expression, which frequently results from AAV-mediated gene delivery, is undesirable as supra-physiological Glut1 is an established risk factor in cancers (42, 43). The ceiling effect associated with the *SLC2A1-DT* NAT that limits Glut1 increase to roughly WT levels resolves this danger. Moreover, use of the lncRNA is expected to stimulate and restore Glut1 only in tissues that express the endogenous transporter physiologically. This property of the lncRNA precludes off-target effects, a second distinct advantage of using the transcript instead of *Glut1* for therapeutic purposes.

While our study has demonstrated the therapeutic value of the *SLC2A1-DT* NAT and clearly shown that the transcript raises *Glut1* expression, it has not revealed the precise mechanism of action of gene induction. Still, some predictions may be made from the existing literature and our own observations. Natural antisense transcripts frequently modulate transcriptional activity by altering chromatin structure. It would therefore not surprise us if such a mechanism underlies the effect of *SLC2A1-DT* on *Glut1* expression. Still, post-transcriptional or even effects on protein translation and/or stability are plausible. Moreover, chromatin modifiers, e.g., the trithorax group of proteins, if they are indeed recruited to the *Glut1* promoter by the lncRNA remain to be identified. Notwithstanding these caveats, which will be addressed in future research, this study has not only presented a potential clinical alternative to the ketogenic diet for Glut1DS but also done so via a unique therapeutic modality – the direct use of a natural antisense transcript.

## Materials and Methods

### Sex as a biological variable

Except for assessing micrencephaly, which is confounded by mixing genders and is only reported for male mice, our experiments did not discriminate between males and females. Note that the trends reported for the micrencephalic phenotype in male animals are similar to those observed in females in our laboratory.

### Primary cell cultures and assessments of human tissue

Human fibroblasts were cultured in M-106 medium (Life Technologies, Carlsbad, CA, USA) supplemented with Low Serum Growth Supplement (LSGS, Cat. # S-003-10), 10% fetal bovine serum (Gibco), and 1% penicillin-streptomycin-glutamine (Fisher Scientific Inc.). Human primary brain microvascular endothelial cells (BECs) were acquired from Cell Biologics (Cat. # H-6023) and grown in complete human endothelial cell medium (Cat. # H1168). To knockdown or overexpress the *SLC2A1-DT* lncRNA, fibroblasts or BECs were seeded in 6-well plates and grown to ∼70% confluency before being transfected. Over-expression of the lncRNA was accomplished by electroporating cells (Nucleofector kit - Cat. # VPD-1001 for fibroblasts; Cat. # VPB-1003 for BECs) with the full-length transcript or truncated versions cloned into the pcDNA3.1 plasmid (Invitrogen). *SLC2A1-DT* knockdown was effected using siRNAs transfected into cells with Lipofectamine 3000 (Invitrogen) according to the manufacturer’s instructions. Seventy-two hours following transfection, all cells were harvested for analysis. Human tissues were obtained from the New York brain bank. Brain tumor tissue was obtained from surgical procedures carried out in the Columbia University department of neurological surgery and banked under IRB AAAJ6163

### Gene reporter assays

A 6kb region upstream of the *SLC2A1* translational start site was cloned into pGL4.10[luc2] vector (Promega) enabling us to set baseline promoter actvity. Effects of the lncRNA on baseline Glut1 promoter activity was assessed in HEK293 cells by co-transfecting pGL4.1-6kb-Glut1 (2.5μg) with a construct containing the full-length NAT in pcDNA3.1 (2.5μg). Assessments were conducted in triplicate and a pSV-βGal (Promega Inc.) used to normalize transfection efficiency. Seventy-two hours following transfection, cells were lysed in reporter lysis buffer (Promega, Madison, WI) and analyzed for luciferase activity (BrightGlo luciferase, Promega Inc.). Luminescence was measured on a GloMax® 20/20 Luminometer (Promega). To assess βGal activity, the cell lysate was incubated in Beta-Glo Reagent (Promega Inc.) at RT for 30min. and luminescence measured as described above.

### FISH analysis

Localization of *SLC2A1-DT* transcripts was determined using the QuantiGene ViewRNA™ ISH Cell Assay Kit (QVC0001, Affymetrix). Briefly, cells were fixed (4% PFA in PBS, 30min. at RT), permeabilized with detergent solution (5min. at RT), and then digested with working protease solution (10min. at RT). Cells were incubated with a target-specific probe for 3hrs., whereas pre-amplifiers, amplifiers, and label probes were incubated for 30min. each. All hybridization steps were carried out at 40°C. Following hybridization, cells were stained with DAPI solution (1min. at RT) and a drop of antifade (Prolong® Gold Antifade Reagent) added to the slide before capturing images on a LEICA TCS SP8 confocal microscope.

### Model mice

Haploinsufficient *Glut1^+/-^* model mice were generated at Columbia University and have been previously described(3). *SLC2A1-DT* transgenic mice were generated for this study by the Columbia University Genetically Modified Mouse Models Core Facility. Briefly, BAC RP11-125O1 containing the human *Glut1* locus was digested with Cla1 and a 79kb fragment harboring *SLC2A1-DT* isolated and introduced into fertilized mouse oocytes. Resulting *SLC2A1-DT^Tg^* mice were interbred with *Glut1^+/-^* animals to produce *Glut1^+/-^;SLC2A1-DT^Tg^* cohorts. Experimental subjects were generated by breeding mutant male mice with WT females. All mice were maintained on a 129 S6/SvEvTac genetic background and mutants identified for experiments by PCR. Littermate controls were used in all experiments.

### Viral vector production and administration

Recombinant scAAV.PhP.eB vectors were produced at the Horae Gene Therapy Core, University of Massachusetts as previously detailed (18, 44). Expression of the lncRNA or eGFP cassettes were driven by a CMV-enhanced chicken-β-actin promoter. Vectors were delivered to PND1 pups at concentrations indicated in the Results section via the retro-orbital sinus.

### Phenotypic evaluations and CSF/blood glucose, lactate measurements

For behavioral outcomes, GraphPad Prism was used to determine sample sizes to detect differences of at least two standard deviations with a power of 80% (*P* < 0.05). Mice were not randomized, but as mutants do not exhibit an overt disease phenotype, it was possible to blind the investigator to the specific cohort being assessed. Motor performance was assessed on an accelerating rotarod (Ugo Basile Inc., Italy) as described in the supplemental information (SI) section. Prior to assessing brain and body size, and investigating CSF/serum glucose levels, mice were fasted overnight. Subsequently they were weighed, blood collected from the tail vein, CSF extracted from the cisterna magna as detailed by us in prior studies (18, 21) and the brain removed and weighed. Glucose concentrations in the CSF and blood were assessed using disposable strips and a Contour Next EZ glucose meter (Bayer Corp., NJ). CSF lactate was measured on an Analox GL5 Analyzer (Analox Instruments, Lunenburg, MA) using a Lactate II reagent kit (GMRD-093, Analox Instruments).

### Quantitative PCR and western blotting

*Glut1* and *SLC2A1-DT* transcript levels were quantified by Q-PCR (see SI for primers). Glut1 protein levels in the various cohorts of mice were assessed by western blot analysis using standard techniques as previously described (18, 21). Briefly, cells or tissues were lysed in lysis buffer containing protease inhibitors (Complete Protease Inhibitor Cocktail tablets, Roche, Indianapolis, IA) and 50μg of total protein resolved by gradient SDS-PAGE (Bio-Rad Labs). Brain parenchymal and vessel fractions were prepared as described in SI. Glut1 rabbit polyclonal (1:5000; Millipore), SMN mouse monoclonal (1:5000; BD Biosciences) and vinculin rabbit monoclonal (1:5000; Abcam) antibodies were probed with HRP-linked goat anti-rabbit (Invitrogen Inc.) and goat anti-mouse (Jackson Immunoresearch) secondary antibodies each diluted 1:10,000 and visualized on a ChemiDoc Imaging System machine (Bio-Rad Labs, Hercules, CA, USA) using the ECL Detection Kit (Cat.# 1705061; Bio-Rad Labs). Band intensities were assessed using ImageJ software (NIH).

### Cerebral microvasculature and neuro-inflammation

To examine the cerebral microvasculature, mice were perfused with 1X PBS and then with 4% PFA. Intact brains were then extracted and further incubated overnight in 4% PFA. Coronal sections (50μm) were cut on a vibratome (LEICA VT1000S) the following day, incubated (1hr., RT) in blocking solution (3% BSA, 0.5% Triton X-100 in PBS) before further incubating them with fluorescein-conjugated Lycopersicon esculentum lectin. The sections were then washed (3X, 20min.) with 1X PBS, mounted with Vectashield (Vector Laboratories) on slides, and overlaid with coverslips for microscopic analysis. Brain capillary density was assessed in 5-month-old mice as previously described(21). Briefly, 20μm stack images of the thalamus were acquired using a LEICA TCS SP8 confocal microscope. Capillary density in thalamic regions involved quantifying the aggregate length of vessels < 6μm in diameter in a 184μm X 184μm area. Three such areas, non-adjacent to one another, in each of at least 4 animals were examined. Images presented are reconstructions of 3D Z-stacks. Note: Total or aggregate length denotes the sum total distance of the stained vessels in the Z-stacks. Gliosis in our subjects was examined and quantified in 15μm stack images of the ventral posteromedial (VPM) thalamic nuclei using antibodies against GFAP and Iba1. Sections were stained and images acquired as described above. Thalamic brain in the region of the VPM nucleus was imaged at a magnification of 20X. Quantification of hypertrophied glial cells was carried out on 63X images. All quantification was carried out by investigators blinded to mouse genotype. Details of antibodies and dilutions employed are supplied in the SI section.

### EEG analysis

Mice were anesthetized using a ketamine (100mg/kg), xylanine (10mg/kg) mixture administered intra-peritoneally and then placed on a stereotaxic frame with a closed-loop heating system to maintain body temperature. After asepsis, the skull was exposed and a small craniotomy (∼0.5mm in diameter) made above the regions of interest. For EEGs, a reference screw was inserted into the skull on top of the cerebellum. EEG recordings were made from two screws on top of the cortex 1mm from midline, 1.5mm anterior to the bregma and 1.5mm posterior to the bregma, respectively. The EEG screws were connected to a PCB board which was soldered with a 5-position pin connector. Implants were secured onto the skull with dental cement (Lang Dental Manufacturing). Animals were given a week to recover before being subjected to recordings. EEG recordings were performed for 24-48 hours (light on at 7:00AM and off at 7:00PM) in a behavioral chamber inside a sound attenuating cubicle (Med Associated Inc.). Subjects were habituated in the chamber for at least 4hrs. before recording began. EEG signals were recorded, bandpass filtered at 0.5-500 Hz, and digitized at 1017 Hz with 32-channel amplifiers (TDT, PZ5 and RZ5D or Neuralynx Digital Lynx 4S). For SWD analysis, FFT of EEG was performed using a 1s sliding window, sequentially shifted by 0.25s increments. Then, the “seizure”-power (19-24 Hz) was calculated to extract SWD events based on a threshold of 2-3 standard deviations as described previously (45). A 19-24 Hz band was selected based on its clear separation from normal brain oscillatory activities, although the primary spectral band of SWDs in mice is around 7Hz, which overlaps with theta oscillations during REM sleep or active periods. Two SWD events were merged into one event if their interval was shorter than 1s. Any SWD event with a duration < 0.5s was removed for analysis. Algorithm-detected SWD events were further reviewed by trained experimenters. Mouse behavior was monitored during recordings using an infrared video camera at 30 frames per second.

### Histology

Hematoxylin/eosin staining experiments were performed on paraffin-embedded 8μm coronal sections (brain) and 4μm longitudinal sections (liver, heart, kidney and muscle) and imaged on an Eclipse 80i Nikon microscope (Nikon, Japan) equipped with a DsRi2 camera (Nikon, Japan).

### Mapping transgene insertion site

To map the insertion site of the transgene harboring the human *Glut1* locus in our engineered mice, we used vectorette PCR. Detailed protocols are presented in the SI section of the manuscript and are modified versions of methods used in previous studies (46, 47).

### RNA-seq and IPA analysis

FASTQ files were aligned to mouse genome assembly GRCm38 using the STAR aligner (48) within the nf-core, rnaseq pipeline (49). Quantification of the aligned data was performed with the default settings of the Salmon tool. After assessing the quality of the data using MultiQC, the resulting counts were analyzed using the limma-voom pipeline (50, 51). Differentially expressed genes (DEGs) were then identified for each pairwise comparison. Note that low and high-dose AAV-lncRNA groups were combined for the purpose of comparisons with control cohorts as we did not observe any significantly altered DEGs between the two lncRNA-treated groups in the RNA-Seq data. To assess for potential toxicity of the lncRNA, we utilized the Tox Analysis function of the Ingenuity Pathway Analysis tool (Qiagen, Inc.), which employs statistical measures to assign z-scores to various toxicological processes predicted to be affected based on the DEGs identified in the RNA-Seq data. In no instance did the analysis show biological process with absolute z-scores greater than 2, which is the cutoff for significant toxicity, for DEGs in the lncRNA-treated subjects.

### Gene ontology analysis

GO analysis was conducted online using the Gene Ontology enrichment analysis & visualization (GORILLA) tool (52). Search parameters were set to identify enriched biological processes at a threshold of *P* < 0.0001. Two unranked lists of genes, a target list of DEGs and a background list of all genes mapped in the RNA-Seq to the Mouse GRCm38.p6 reference genome were queried to identify pathways significantly altered in mutants.

### Statistics

For data that was normally distributed, the unpaired 2-tailed Student’s *t* test or 1-way ANOVA followed by Tukey’s post-hoc comparison, where indicated, were used to compare means for statistical differences. If the data was not found to fit a Gaussian distribution, the Mann-Whitney test or Kruskal-Wallis test followed by Dunn’s multiple comparison test were used to compare ranks. Data in the manuscript are represented as mean ± SEM. *P* < 0.05 was considered significant. Statistical analyses were performed with GraphPad Prism v9.0 (GraphPad Software).

### Study approval

All animal procedures adhered to protocols described in the *Guide for the Care & Use of Lab Animals* (National Academic Press, 2011) and were approved by Columbia University’s IACUC. The subjects of this study were randomly selected, 129 S6/SvEvTac, male and female mice housed in a controlled environment on a 12hr. light/dark cycle with food and water.

## Data availability

Underlying values associated with the data presented in the study may be requested and will be made available by the lead contact, Umrao R. Monani.

## Acknowledgments

We are grateful to members of the Giblin Laboratories for their comments and suggestions. Work in our laboratories was funded by the Glut1 Deficiency Foundation (U.R.M., D.C.D), The University of Pennsylvania Orphan Disease Center (U.R.M.), the Hope for Children Research Foundation (U.R.M., D.C.D), AFM-France (U.R.M.), the Crofoot family foundation, Sarepta Therapeutics Inc. and NIH (R03 NS128211). Y.P. acknowledges support from the Columbia Precision Medicine Initiative. P.L.F. received funding from NINDS (R01 NS086736, R01 NS117745 and R01 NS124854), and P.C. and J.N.B. acknowledge support from the Columbia University Cancer Center and from NCI (P30CA013696).

## Author contributions

M.T. planned and performed most of the experiments described here. S.T. and Y.P. carried out the EEG analyses. P.L.F. assisted with the histological analysis of the mice, while P.C. and J.N.B. acquired the brain tumor tissue for this study. A.Y.K. was involved in animal husbandry and genotyping mice for the study, and K.A. analyzed the RNA-Seq data bioinformatically. D.C.D. provided intellectual input and helped prepare the manuscript. U.R.M. conceptualized the experiments, directed the project, analyzed and interpreted the data. U.R.M. and M.T. wrote the manuscript with input from all authors.

## Supplemental Information

### Rotarod test

The rotarod test was administered by subjecting mice to a training period of 5mins. on an accelerating rotarod from 0-40rpm (Ugo Basile Inc., Italy) three times a day for four consecutive days. Performance on day 5 was assessed at a setting of 25rpm. The rotarod was cleaned and dried prior to testing. Latency to fall off the rotating rod was recorded and the experiment terminated if a mouse surpassed 1000s.

### Preparation of brain parenchymal and vessel fractions

Following euthanasia and perfusion with 1X PBS, mouse brains were extracted and gently homogenized using a Dounce-type glass homogenizer in 1ml PBS. The resulting extract was centrifuged at 1000g for 5min., the pellet re-suspended once again in 1ml 1X PBS and the centrifugation step repeated. The supernatant was then discarded and the pellet re-suspended (1ml 18% dextran solution in PBS). This suspension was centrifuged (10,000g for 1min.), the pellet retrieved, and the supernatant containing the neuropil fraction transferred to new tube. The pellet was once again re-suspended (1ml 18% dextran in PBS) and the centrifugation repeated. This process was repeated a third time and the pellets containing the vessel fractions were pooled and stored at -80°C until use. The supernatant fractions – containing the neuropil – were similarly combined and stored for analysis.

### Vectorette PCR

To map the insertion site of the transgene bearing the human Glut1 locus, we used a vectorette PCR protocol previously described in references 46 and 47. In brief, genomic DNA from transgenic mice was digested individually with each of the following enzymes – RsaI, HincII, HpaI and ScaI – to create corresponding libraries. Subsequently, a linker (vectorette) consisting of 52bp strands that exhibit complementarity at their ends but not in a central 29bp central region was ligated onto blunt ends generated by the various restriction enzymes. Such libraries were then subjected to an initial round of PCR using a transgene specific primer and a vectorette primer (UVP) that binds to sequence in the “bubble” formed within the vectorette. Amplification products were then used as template for a nested PCR using the vectorette primer and a second transgene specific primer. This increases specificity of the PCR and generally resulted in one distinct product. In instances where the nested PCR did not result in a single distinct PCR product, a double-nested PCR was carried out using amplification products from the nested PCR as template. The final PCR products were then sequenced, junction fragments determined and sequence in these fragments corresponding to the mouse genome mapped onto the murine reference genome (GRCM39) to identify the precise insertion site of the transgene.

### Quantitative PCR

RNA for quantitative RT-PCR was extracted from relevant tissue or cells using Trizol (Life Technologies, Carlsbad, CA) according to the manufacturer’s instructions. Following treatment with DNAse, cDNA was synthesized using the RevertAid First Strand cDNA Synthesis Kit (Thermo Scientific). QPCRs were carried out on a CFX96 Real-Time Systems PCR analyzer (Bio-Rad Labs) using the SsoAdvanced Universal SYBR Green Supermix (Bio-Rad Labs). Relative expression of the *Glut1* gene and the *SLC2A1-DT* lncRNA was calculated using the ΔΔCT method. Absolute amounts of *Glut1* and *SLC2A1-DT* RNAs in brain tumor samples were measured following construction of a standard curve. These standard curves were constructed using serial 10-fold dilutions of purified plasmid containing the target sequence. Linear regression was used to calculate the slope and intercept for the standard curve.

### Key Reagents

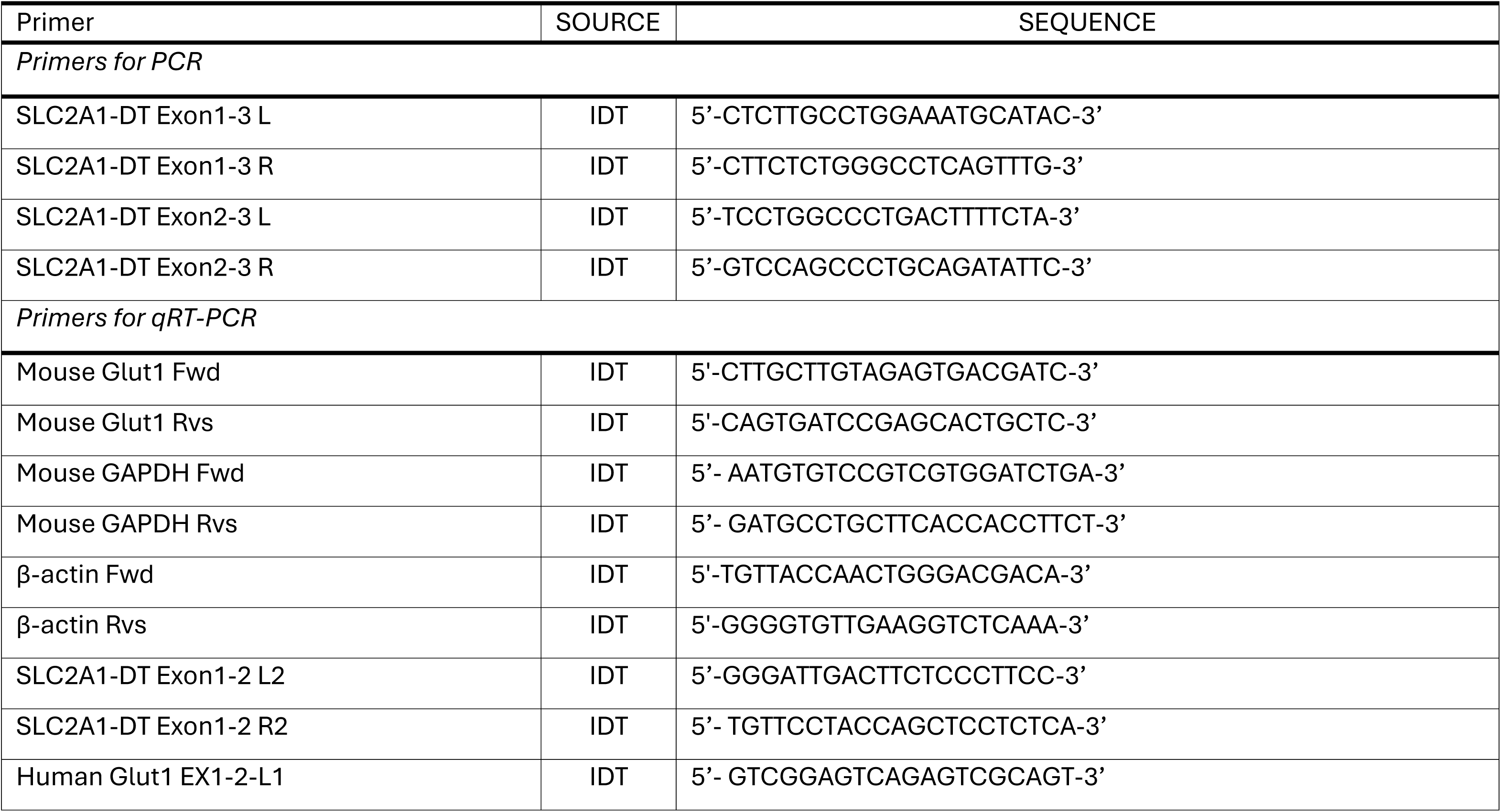
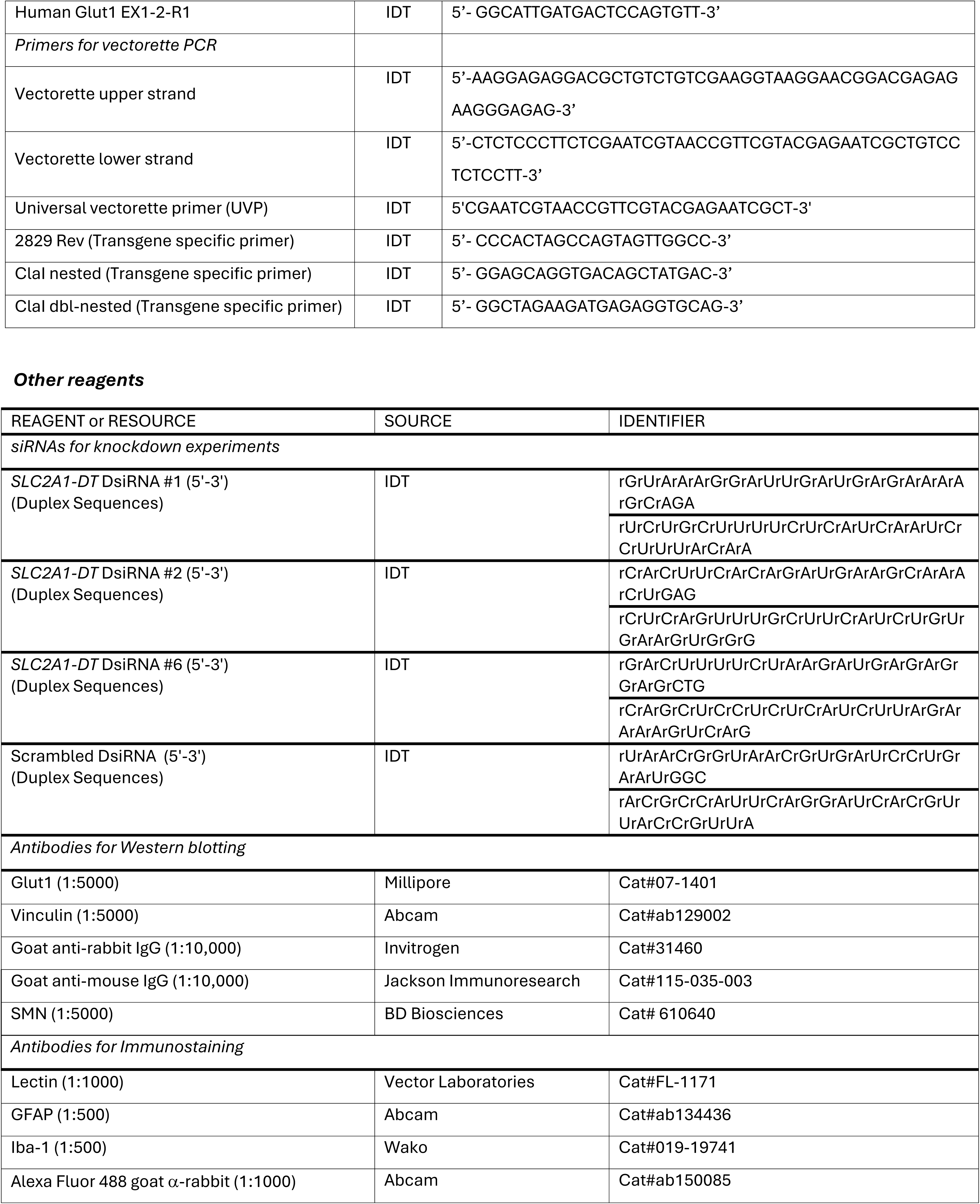
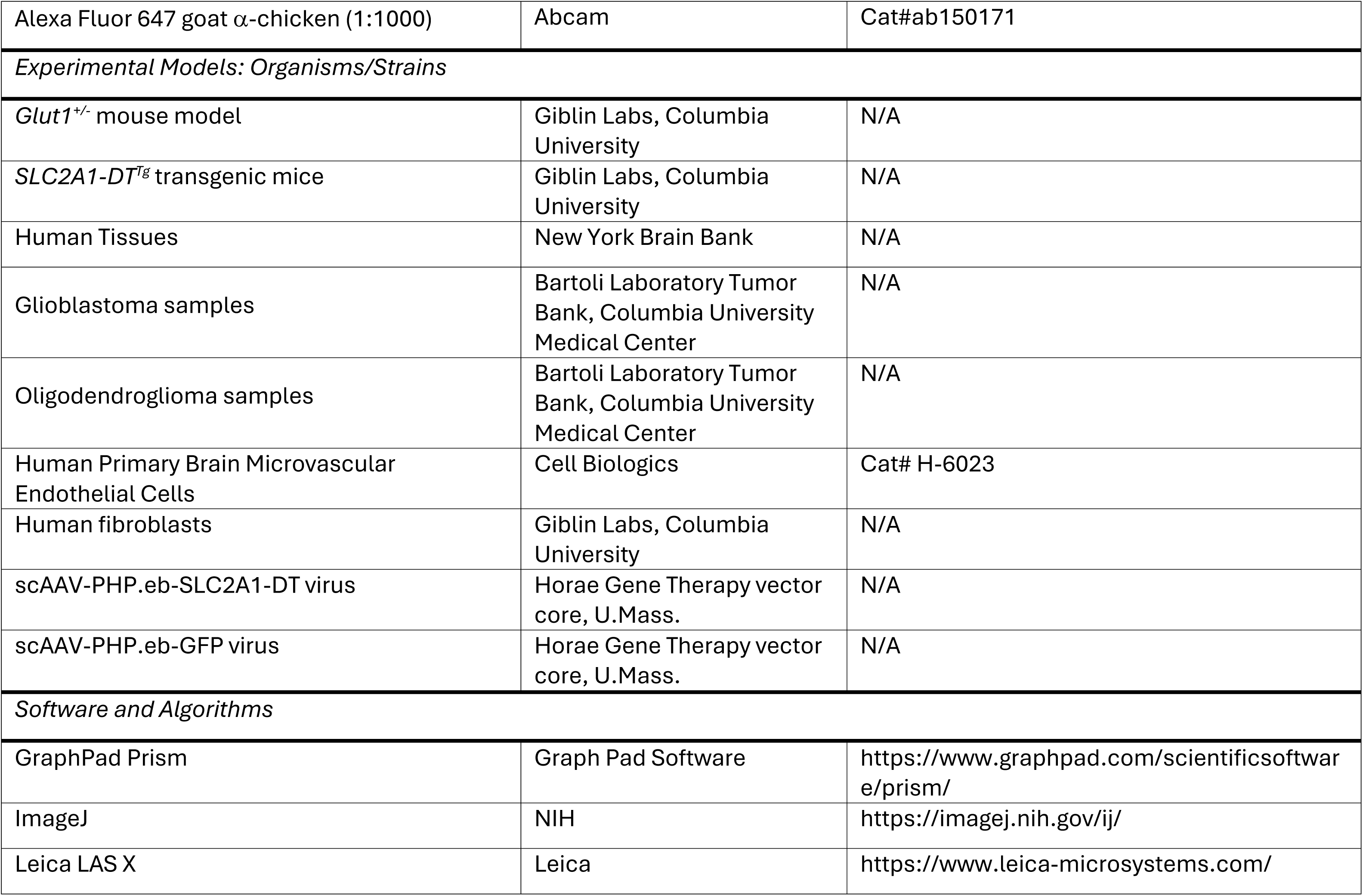

---

**Supplemental Fig. 1.**
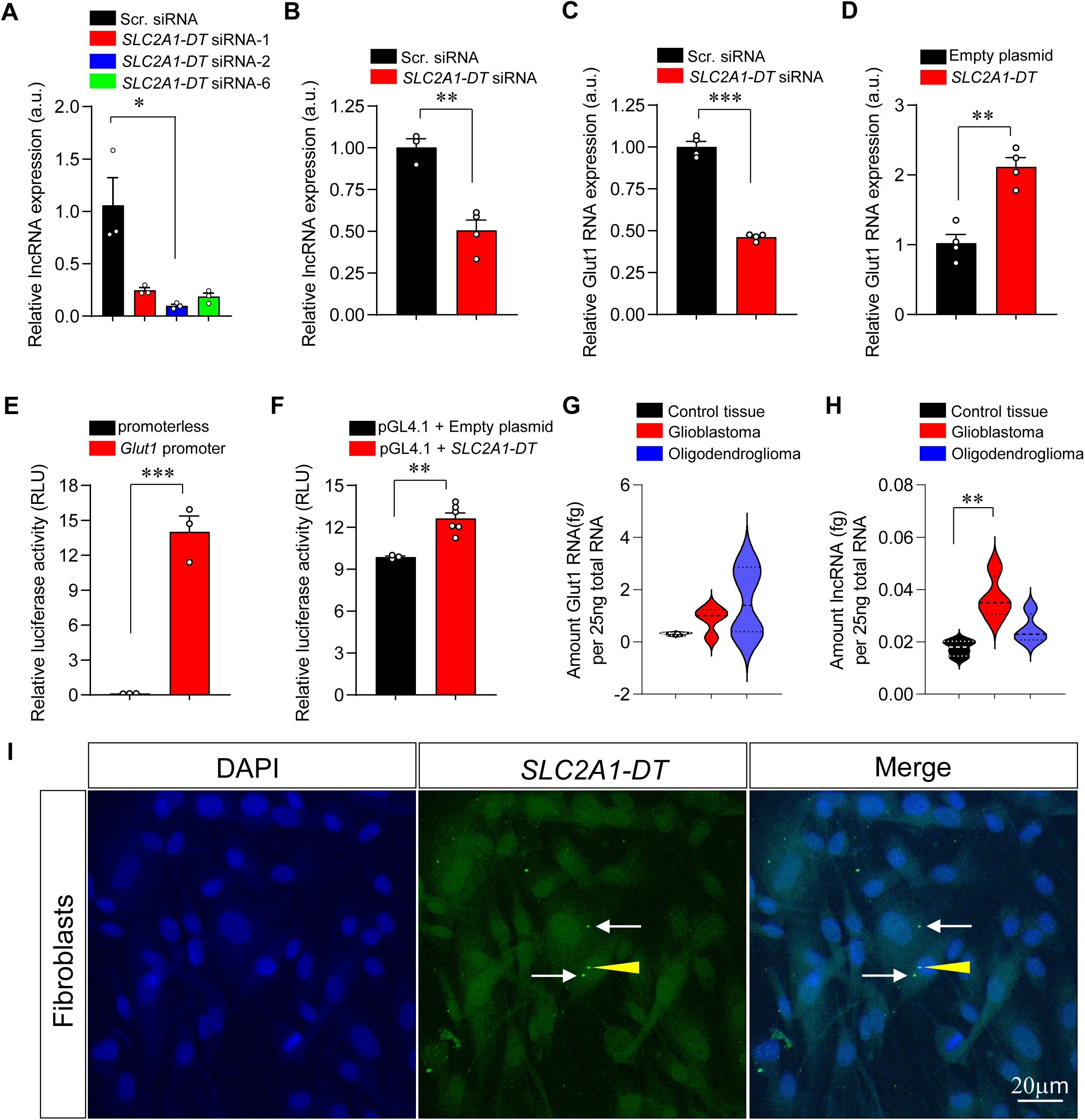
The *SLC2A1-DT* lncRNA influences *Glut1* expression. **(A)** lncRNA expression in human fibroblasts is reduced by siRNAs. *, *P* < 0.05, Kruskal-Wallis test, n = 3 experiments for each condition. **(B)** Quantified results of *SLC2A1-DT* knockdown by siRNA #2 in human brain endothelial cells (BECs). **, *P* < 0.01, *t* test, n ≥ 3 replicates. **(C)** Knockdown of *SLC2A1-DT* suppressed *Glut1* expression significantly in human BECs. ***, *P* < 0.001, *t* test, n = 4 replicates. **(D)** Over-expressing *SLC2A1-DT* raised Glut1 RNA levels in human BECs. **, *P* < 0.01, *t* test, n = 4 replicates. **(E)** A human *Glut1* promoter but not promoterless construct drives robust reporter expression. ***, *P* < 0.001, *t* test, n = 3 replicates. **(F)** Graph depicting increased reporter activity in HEK293 cells co-transfected with a construct expressing *SLC2A1-DT*. **, *P* < 0.001, *t* test, n ≥ 3 replicates each. An increase in **(G)** Glut1 expression in human brain tumor samples is mirrored by a similar increase **(H)** in lncRNA levels. **, *P* < 0.01, one-way ANOVA, n ≥ 3 samples in each instance. **(I)** Representative photomicrographs of fibroblasts illustrating the localization of the *SLC2A1-DT* lncRNA. Arrows depict lncRNA puncta in the cell soma whereas the arrowhead highlights the transcript in the nucleus.

**Supplemental Fig. 2.**
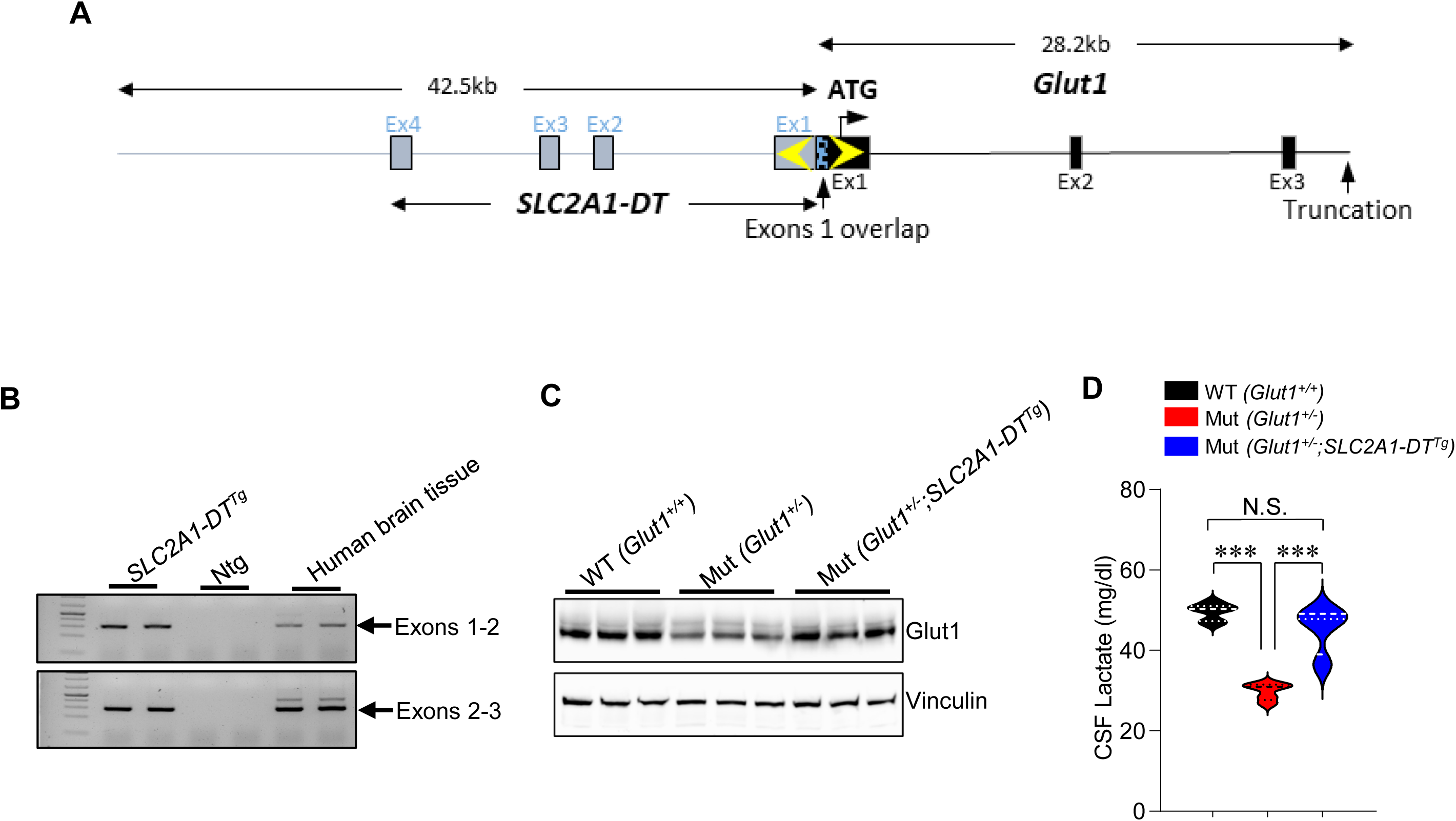
Transgenic expression and disease amelioration by the *SLC2A1-DT* lncRNA. **(A)** Cartoon depicting the structure of the transgene bearing the *SLC2A1-DT* lncRNA. Note: Truncation of the human *Glut1* gene in the transgene renders *Glut1* non-functional. **(B)** Representative gel following qualitative PCR depicts expression of the lncRNA transcript in transgenic mice but not in littermates devoid of the transgene. Bands correspond to gene segments amplified using primers lying in exons 1 and 2 or in exons 2 and 3. **(C)** Western blot illustrating an increase in murine Glut1 protein in transgenic mice expressing the *SLC2A1-DT* lncRNA transgene. **(D)** Graph depicting normalization of CSF lactate levels in Glut1DS mutant mice expressing the *SLC2A1-DT* lncRNA transgene. ***, *P* < 0.001, one-way ANOVA, n ≥ 4 mice of each cohort.

**Supplemental Fig. 3.**
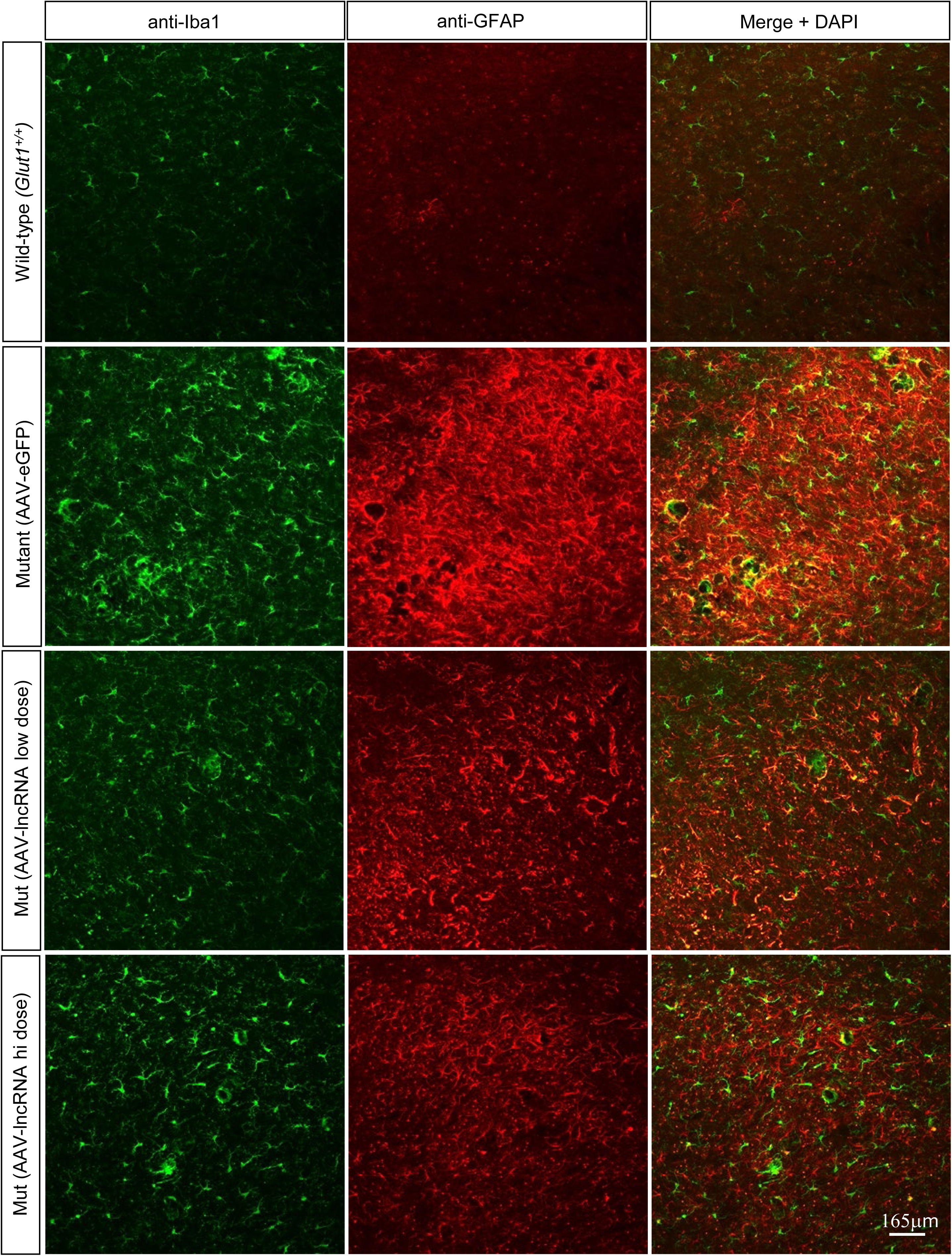
The AAV-*SLC2A1-DT* lncRNA mitigates the neuroinflammatory response in Glut1DS model mice. **(A)** Representative thalamic sections, imaged at low magnification, from 4 – 5-month-old controls and mutants treated either with AAV-eGFP or AAV-lncRNA and stained with antibodies against Iba1 and GFAP to assess gliosis in the various cohorts of mice. Note profound neuroinflammation in mutants treated with the AAV-eGFP construct; cerebral gliosis is reduced in Glut1DS mice administered the lncRNA-containing AAV vector.

**Supplemental Fig. 4.**
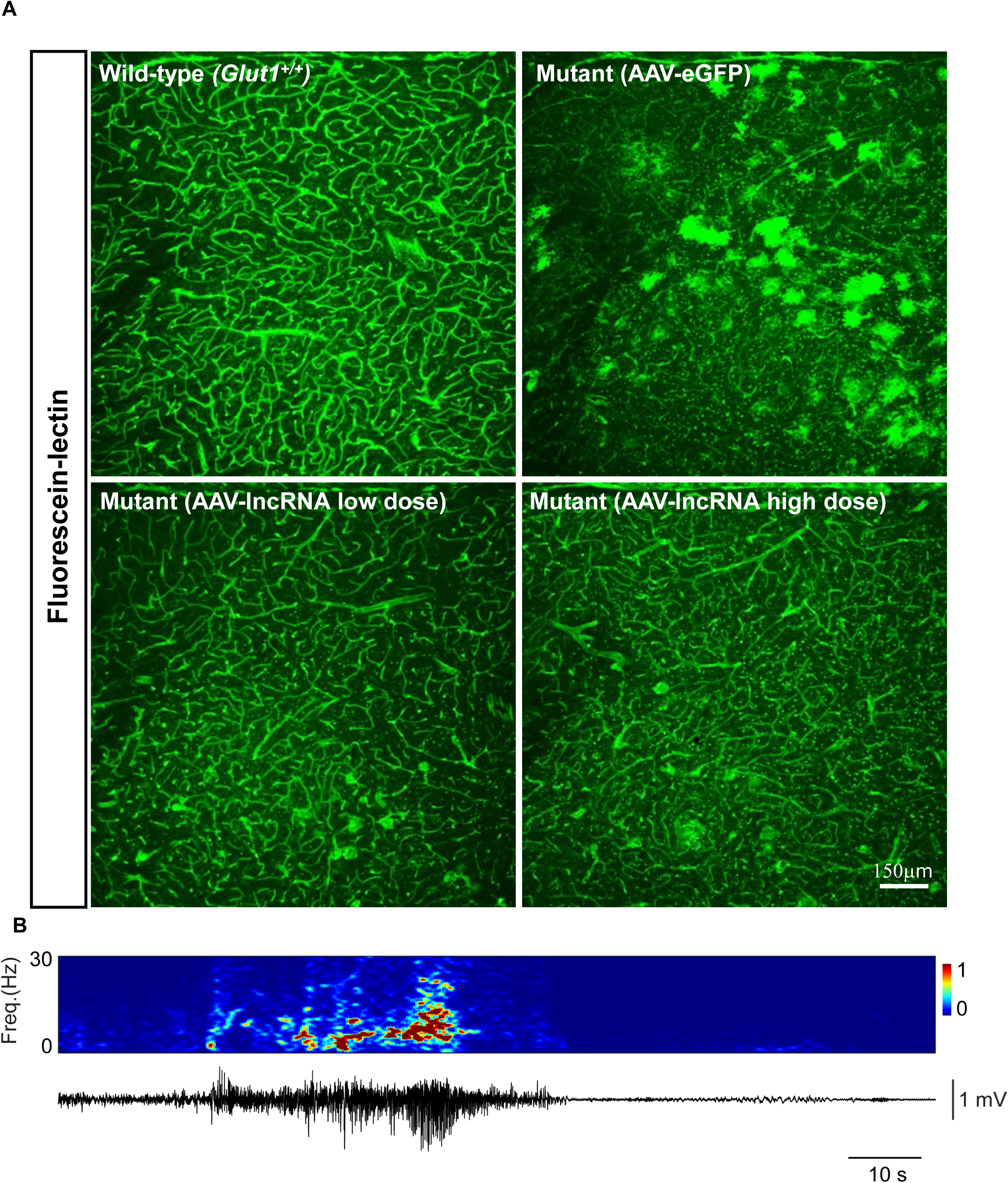
The AAV-*SLC2A1-DT* lncRNA mitigates cerebral pathology in Glut1DS model mice. **(A)** Representative thalamic sections, imaged at low magnification, from 4 – 5-month-old control and mutants treated either with AAV-eGFP or AAV-lncRNA and stained with fluorescein-linked lectin to reveal the brain microvasculature. Note fewer brain capillaries in the AAV-eGFP-treated individual and relative normalization of brain angiogenesis in the mutants treated with the lncRNA; intense fluorescence in mutants injected with AAV-eGFP derives from reporter expressed in cells of the neuropil **(B)** Representative EEG spectrogram (at 0-25Hz) and corresponding trace below it depicting a convulsive seizure in a Glut1DS mutant delivered the control (AAV-eGFP) vector. These were eliminated in mutants treated with the AAV-lncRNA construct.

**Supplemental Fig. 5.**
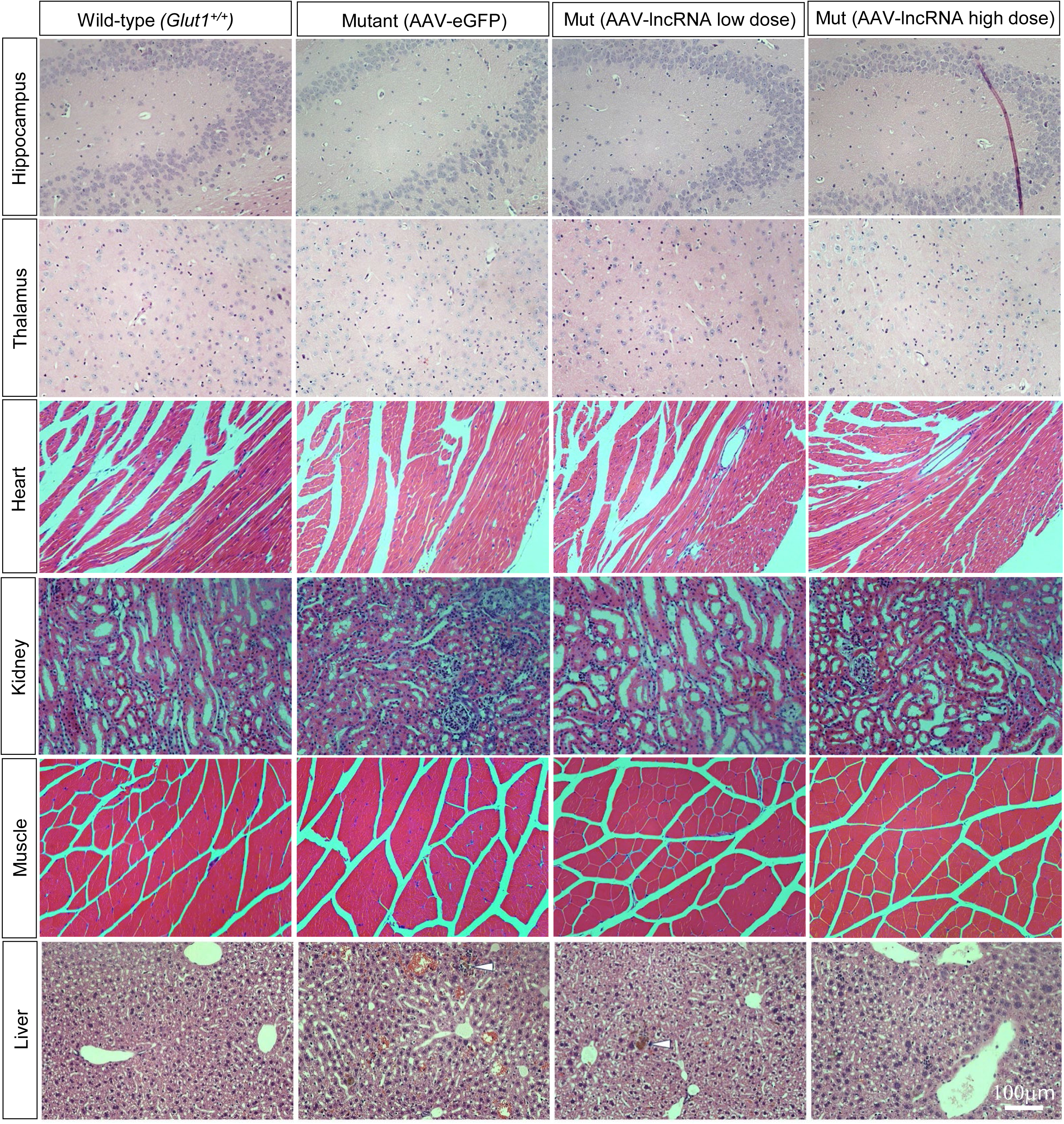
Early administration of the AAV-*SLC2A1-DT* lncRNA does not trigger major organ pathology in model mice. Hematoxylin/eosin histochemistry of the major organs of the body at 12 months of age depicts normal cellular morphology in most tissues of mutants treated with AAV-lncRNA. Cholestatic bile pigment in liver of lncRNA treated mice (arrowhead) was also observed in AAV-eGFP-treated mutants

**Supplemental Table 1.** DEGs: AAV-lncRNA versus WT mice.

|  | baseMean | log2FoldCI | lfcSE | stat | pvalue | padj |
| --- | --- | --- | --- | --- | --- | --- |
| Slc2a1 | 1765.767 | -0.65404 | 0.055821 | -11.7167 | 1.05E-31 | 1.39E-27 |
| Olfr1438-p | 131.585 | -0.58338 | 0.120167 | -4.85477 | 1.21E-06 | 0.002294 |
| Pou6f2 | 160.0661 | -0.48481 | 0.119434 | -4.05927 | 4.92E-05 | 0.022561 |
| Haus7 | 169.2629 | -0.46837 | 0.104937 | -4.46329 | 8.07E-06 | 0.008962 |
| Prr5l | 416.5238 | -0.39345 | 0.094846 | -4.14835 | 3.35E-05 | 0.018593 |
| Gm29216 | 114709.7 | -0.35804 | 0.07288 | -4.9127 | 8.98E-07 | 0.001995 |
| Gcnt4 | 461.8583 | -0.33452 | 0.081197 | -4.11983 | 3.79E-05 | 0.019431 |
| Six3 | 635.0703 | -0.29859 | 0.070479 | -4.23662 | 2.27E-05 | 0.014398 |
| Npnt | 914.1626 | -0.28324 | 0.073458 | -3.85581 | 0.000115 | 0.037488 |
| Kcnab3 | 2901.082 | -0.25525 | 0.057546 | -4.43562 | 9.18E-06 | 0.009365 |
| Arglu1 | 7798.597 | -0.25304 | 0.054578 | -4.63634 | 3.55E-06 | 0.00525 |
| Fryl | 3034.816 | -0.24331 | 0.060863 | -3.9976 | 6.40E-05 | 0.026645 |
| mt-Co2 | 293108 | -0.24035 | 0.04228 | -5.68462 | 1.31E-08 | 5.82E-05 |
| Kifc3 | 1227.172 | -0.2374 | 0.057399 | -4.13605 | 3.53E-05 | 0.018833 |
| Pdcd6ip | 3719.243 | -0.23113 | 0.060552 | -3.81702 | 0.000135 | 0.041857 |
| Uqcrq | 1343.812 | -0.2246 | 0.049761 | -4.5136 | 6.37E-06 | 0.008493 |
| Samd5 | 1243.667 | -0.20232 | 0.049931 | -4.05194 | 5.08E-05 | 0.022561 |
| Mbp | 74949.69 | -0.1997 | 0.038061 | -5.24683 | 1.55E-07 | 0.000515 |
| Cpsf3 | 1513.511 | -0.19265 | 0.046899 | -4.10783 | 3.99E-05 | 0.019711 |
| Prpf38b | 3352.428 | -0.16712 | 0.04403 | -3.79568 | 0.000147 | 0.04459 |
| Kcnip4 | 3382.74 | -0.15933 | 0.041121 | -3.87467 | 0.000107 | 0.035568 |
| Scrn1 | 10551.3 | -0.13568 | 0.035484 | -3.82381 | 0.000131 | 0.04169 |
| Ypel3 | 4559.617 | 0.12326 | 0.031784 | 3.878042 | 0.000105 | 0.035568 |
| Rps3a2 | 5248.874 | 0.124058 | 0.031489 | 3.939712 | 8.16E-05 | 0.031111 |
| Stx7 | 4095.748 | 0.148158 | 0.037627 | 3.937571 | 8.23E-05 | 0.031111 |
| Rps15a | 3790.927 | 0.154516 | 0.035647 | 4.334652 | 1.46E-05 | 0.010807 |
| Zhx3 | 2645.199 | 0.158201 | 0.036808 | 4.298032 | 1.72E-05 | 0.011481 |
| Zcchc24 | 3604.61 | 0.188238 | 0.042816 | 4.396437 | 1.10E-05 | 0.009775 |
| C1ql3 | 1501.862 | 0.221828 | 0.05696 | 3.894445 | 9.84E-05 | 0.035446 |
| Ftl1 | 4395.846 | 0.22645 | 0.038021 | 5.955984 | 2.59E-09 | 1.72E-05 |
| Acot11 | 2199.16 | 0.240761 | 0.053664 | 4.48642 | 7.24E-06 | 0.008774 |
| Sap30bp | 1312.191 | 0.271324 | 0.062912 | 4.312722 | 1.61E-05 | 0.011309 |
| Gfap | 7683.471 | 0.277958 | 0.066596 | 4.173796 | 3.00E-05 | 0.017355 |
| Phactr4 | 886.7514 | 0.312222 | 0.066131 | 4.721245 | 2.34E-06 | 0.003904 |
| Fn1 | 2031.682 | 0.325211 | 0.073566 | 4.420675 | 9.84E-06 | 0.009365 |
| Cdyl2 | 200.2533 | 0.336817 | 0.084075 | 4.006147 | 6.17E-05 | 0.026528 |
| Ahnak | 1338.068 | 0.347921 | 0.079594 | 4.371208 | 1.24E-05 | 0.009961 |
| Ccn3 | 1573.852 | 0.396399 | 0.09426 | 4.205396 | 2.61E-05 | 0.015786 |
| Yrdc | 587.8215 | 0.445156 | 0.089911 | 4.951086 | 7.38E-07 | 0.001967 |
| Slc6a20a | 922.9108 | 0.449954 | 0.114418 | 3.932542 | 8.41E-05 | 0.031111 |
| Ccdc36 | 306.6696 | 0.488591 | 0.123298 | 3.962683 | 7.41E-05 | 0.029926 |
| Aebp1 | 559.0145 | 0.498325 | 0.122374 | 4.072164 | 4.66E-05 | 0.022166 |
| Dynlt1-ps1 | 163.1502 | 0.528201 | 0.121006 | 4.365076 | 1.27E-05 | 0.009961 |
| Mrc1 | 116.9699 | 0.731557 | 0.188796 | 3.874865 | 0.000107 | 0.035568 |

Note: Only DEGs with Padj values < 0.05 are listed; down-regulated and up-regulated genes in AAV-lncRNA mutants in blue and red text respectively. Also note: Mutants administered low and high dose of AAV-lncRNA were combined into one group to carry out this analysis.

**Supplemental Table 2.**
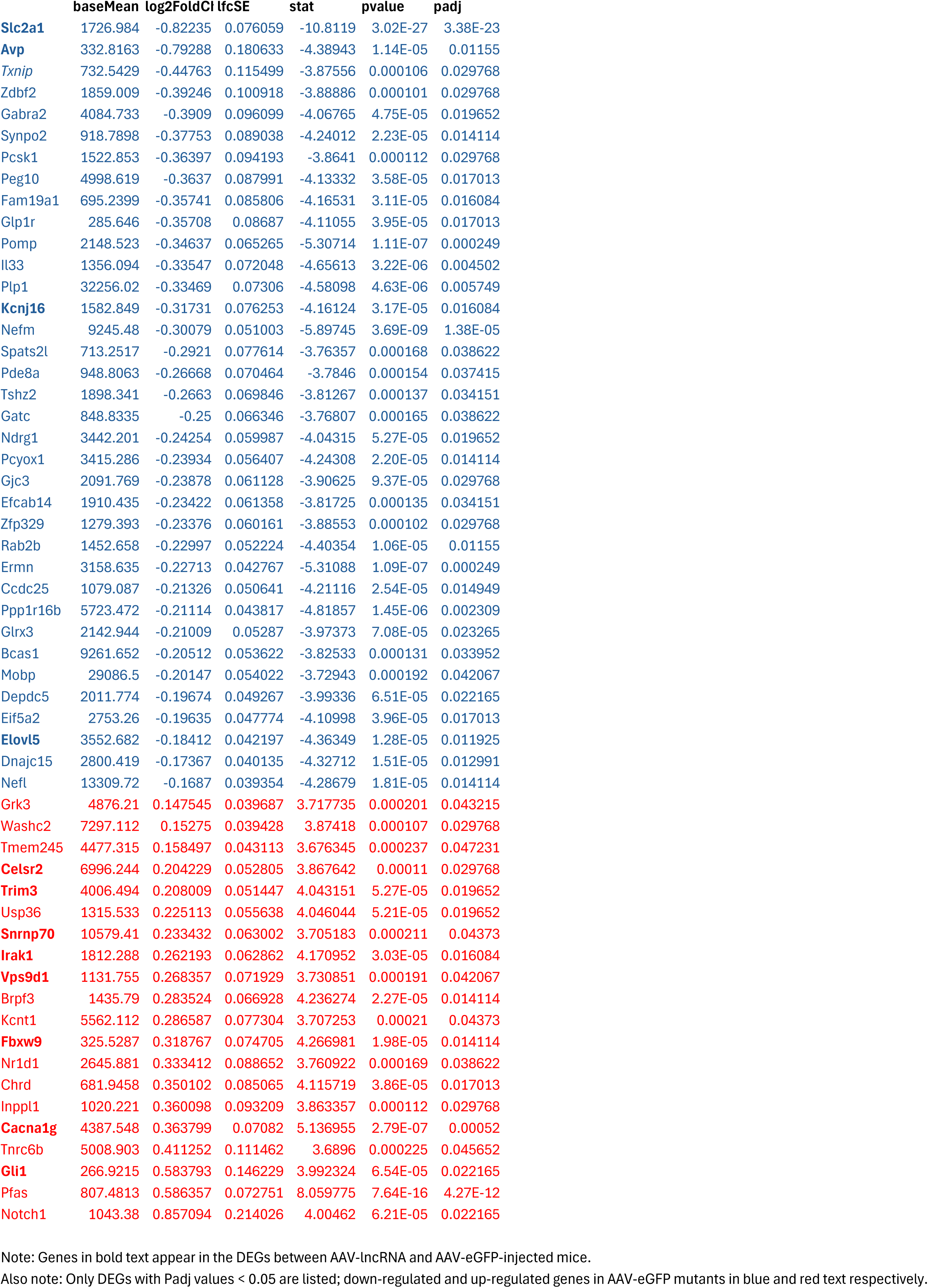
DEGs: AAV-eGFP versus WT mice.

**Supplemental Table 3.** DEGs: AAV-lncRNA versus AAV-eGFP mice.

|  | baseMean | log2FoldCh | lfcSE | stat | pvalue | padj |
| --- | --- | --- | --- | --- | --- | --- |
| Gm15542 | 113.0132 | -1.003685 | 0.228155 | -4.399129 | 1.09E-05 | 0.001338 |
| Muc3a | 90.95299 | -0.821982 | 0.198295 | -4.145254 | 3.39E-05 | 0.00305 |
| Rtbdn | 110.4235 | -0.796537 | 0.193023 | -4.126638 | 3.68E-05 | 0.003167 |
| **Gli1** | 274.6103 | -0.729834 | 0.169231 | -4.312643 | 1.61E-05 | 0.001756 |
| Robo3 | 479.8148 | -0.728554 | 0.169528 | -4.297541 | 1.73E-05 | 0.001826 |
| Gm49405 | 167.9824 | -0.66359 | 0.153934 | -4.310865 | 1.63E-05 | 0.001756 |
| Olfr152 | 354.9549 | -0.656299 | 0.128265 | -5.116728 | 3.11E-07 | 8.33E-05 |
| Fdxr | 317.2011 | -0.653136 | 0.121595 | -5.371396 | 7.81E-08 | 2.91E-05 |
| Rgs11 | 645.549 | -0.611366 | 0.107614 | -5.681118 | 1.34E-08 | 7.17E-06 |
| Mri1 | 188.0286 | -0.590311 | 0.143906 | -4.102068 | 4.09E-05 | 0.003405 |
| Miip | 210.5351 | -0.58755 | 0.14791 | -3.972356 | 7.12E-05 | 0.004862 |
| Mkx | 398.7035 | -0.585729 | 0.133767 | -4.378717 | 1.19E-05 | 0.001435 |
| Cdc25b | 473.4333 | -0.579897 | 0.105447 | -5.499418 | 3.81E-08 | 1.76E-05 |
| Spaca6 | 1543.639 | -0.57745 | 0.096085 | -6.009769 | 1.86E-09 | 1.78E-06 |
| Gm5373 | 616.1565 | -0.561552 | 0.127549 | -4.402652 | 1.07E-05 | 0.001338 |
| Actn2 | 2672.208 | -0.561408 | 0.095169 | -5.899066 | 3.66E-09 | 2.72E-06 |
| Sema4a | 1160.425 | -0.546233 | 0.136923 | -3.98936 | 6.63E-05 | 0.004629 |
| Ulk3 | 599.0907 | -0.538368 | 0.093282 | -5.771411 | 7.86E-09 | 4.58E-06 |
| Cc2d1b | 898.3271 | -0.517515 | 0.094763 | -5.461151 | 4.73E-08 | 2.11E-05 |
| Vwa5b2 | 1471.908 | -0.509278 | 0.120595 | -4.223054 | 2.41E-05 | 0.002356 |
| Zfp46 | 1226.261 | -0.509266 | 0.123011 | -4.140014 | 3.47E-05 | 0.00308 |
| Rgl2 | 991.8693 | -0.498337 | 0.076954 | -6.475807 | 9.43E-11 | 1.80E-07 |
| Phka2 | 1006.096 | -0.494013 | 0.119094 | -4.148083 | 3.35E-05 | 0.003033 |
| Ssh3 | 1123.758 | -0.487917 | 0.09435 | -5.171373 | 2.32E-07 | 7.07E-05 |
| Ctu2 | 295.6667 | -0.48588 | 0.108287 | -4.486965 | 7.22E-06 | 0.000986 |
| Dennd6b | 1536.518 | -0.479309 | 0.072554 | -6.606269 | 3.94E-11 | 1.06E-07 |
| Prpf40b | 2032.096 | -0.478177 | 0.069847 | -6.846018 | 7.59E-12 | 5.08E-08 |
| Vmn2r115 | 200.8129 | -0.477642 | 0.110957 | -4.30476 | 1.67E-05 | 0.001791 |
| Carmil3 | 1781.468 | -0.464715 | 0.07948 | -5.84696 | 5.01E-09 | 3.35E-06 |
| Coro6 | 1854.557 | -0.461873 | 0.089902 | -5.137503 | 2.78E-07 | 8.04E-05 |
| Acp6 | 536.6184 | -0.454734 | 0.080453 | -5.652183 | 1.58E-08 | 7.86E-06 |
| Myo19 | 371.3866 | -0.445551 | 0.111183 | -4.007375 | 6.14E-05 | 0.004501 |
| Srrm2 | 25544.44 | -0.445499 | 0.067064 | -6.64285 | 3.08E-11 | 1.03E-07 |
| Prss12 | 472.938 | -0.443455 | 0.111125 | -3.990606 | 6.59E-05 | 0.004629 |
| Zfp280c | 948.5307 | -0.432166 | 0.087287 | -4.951114 | 7.38E-07 | 0.00017 |
| Dmpk | 714.1947 | -0.42573 | 0.101073 | -4.212114 | 2.53E-05 | 0.002404 |
| **Cacna1g** | 4455.051 | -0.424119 | 0.073051 | -5.805786 | 6.41E-09 | 4.08E-06 |
| Snhg11 | 72685.76 | -0.416515 | 0.078615 | -5.298181 | 1.17E-07 | 4.12E-05 |
| **Fbxw9** | 319.651 | -0.410046 | 0.102543 | -3.998765 | 6.37E-05 | 0.004584 |
| Arglu1 | 7561.853 | -0.40937 | 0.053835 | -7.604156 | 2.87E-14 | 3.84E-10 |
| Mbd6 | 1891.449 | -0.40368 | 0.064163 | -6.29145 | 3.15E-10 | 4.61E-07 |
| Leng8 | 10146.88 | -0.403455 | 0.060435 | -6.675854 | 2.46E-11 | 1.03E-07 |
| Adamts3 | 518.2058 | -0.400834 | 0.088141 | -4.54764 | 5.43E-06 | 0.000776 |
| **Vps9d1** | 1084.906 | -0.399344 | 0.073933 | -5.401424 | 6.61E-08 | 2.60E-05 |
| Tspoap1 | 6499.033 | -0.396998 | 0.083335 | -4.763876 | 1.90E-06 | 0.000371 |
| Usp35 | 527.5444 | -0.394541 | 0.09111 | -4.330393 | 1.49E-05 | 0.001689 |
| Tctn1 | 822.3524 | -0.383289 | 0.091165 | -4.204342 | 2.62E-05 | 0.002452 |
| Lrrc45 | 1204.493 | -0.383185 | 0.081225 | -4.717589 | 2.39E-06 | 0.00043 |
| Anks3 | 994.3823 | -0.38273 | 0.090701 | -4.21969 | 2.45E-05 | 0.00237 |
| Celsr3 | 4540.687 | -0.376787 | 0.062693 | -6.010078 | 1.85E-09 | 1.78E-06 |
| Ncapd3 | 1048.743 | -0.372166 | 0.079819 | -4.662595 | 3.12E-06 | 0.000523 |
| Doc2a | 801.2738 | -0.371989 | 0.088182 | -4.218452 | 2.46E-05 | 0.00237 |
| Odf2 | 2864.021 | -0.371512 | 0.068939 | -5.389016 | 7.08E-08 | 2.71E-05 |
| Syt3 | 1134.725 | -0.371299 | 0.092728 | -4.004162 | 6.22E-05 | 0.004529 |
| Cacna1h | 1769.327 | -0.371027 | 0.064147 | -5.78406 | 7.29E-09 | 4.44E-06 |
| Sema4g | 1627.024 | -0.367132 | 0.079256 | -4.632254 | 3.62E-06 | 0.000588 |
| Nav2 | 2560.988 | -0.363121 | 0.072504 | -5.008292 | 5.49E-07 | 0.000134 |
| Dot1l | 3548.713 | -0.360223 | 0.090273 | -3.990389 | 6.60E-05 | 0.004629 |
| Lars | 1580.709 | -0.355771 | 0.074757 | -4.759059 | 1.94E-06 | 0.000372 |
| Rsrp1 | 8549.943 | -0.350982 | 0.074246 | -4.727318 | 2.28E-06 | 0.000417 |
| Rcor3 | 1087.155 | -0.345224 | 0.083197 | -4.149481 | 3.33E-05 | 0.003033 |
| Ankrd16 | 997.0427 | -0.344371 | 0.085394 | -4.03271 | 5.51E-05 | 0.004202 |
| Clasrp | 2094.751 | -0.344152 | 0.060735 | -5.666414 | 1.46E-08 | 7.51E-06 |
| Sec61a2 | 2259.321 | -0.34402 | 0.081087 | -4.242631 | 2.21E-05 | 0.002228 |
| Atp11b | 4591.118 | -0.343744 | 0.058214 | -5.904802 | 3.53E-09 | 2.72E-06 |
| Rps6ka4 | 1077.71 | -0.341674 | 0.074673 | -4.575603 | 4.75E-06 | 0.000723 |
| Plekhn1 | 1393.598 | -0.341627 | 0.065892 | -5.184639 | 2.16E-07 | 6.74E-05 |
| Srrm1 | 7243.343 | -0.340593 | 0.059745 | -5.700832 | 1.19E-08 | 6.65E-06 |
| Hdx | 648.9536 | -0.339138 | 0.08502 | -3.988916 | 6.64E-05 | 0.004629 |
| Scaf4 | 1835.765 | -0.338064 | 0.064231 | -5.263264 | 1.42E-07 | 4.74E-05 |
| Aifm3 | 3624.71 | -0.333225 | 0.081146 | -4.106485 | 4.02E-05 | 0.003383 |
| Map4k2 | 2653.318 | -0.328428 | 0.079363 | -4.138308 | 3.50E-05 | 0.003082 |
| Cacna1a | 9096.929 | -0.321791 | 0.081135 | -3.966119 | 7.31E-05 | 0.004915 |
| **Trim3** | 3935.082 | -0.321121 | 0.068288 | -4.702442 | 2.57E-06 | 0.000441 |
| Nsmaf | 1926.8 | -0.318427 | 0.062011 | -5.135007 | 2.82E-07 | 8.04E-05 |
| **Snrnp70** | 10139.34 | -0.317372 | 0.053248 | -5.960227 | 2.52E-09 | 2.25E-06 |
| Stk38 | 1709.347 | -0.315758 | 0.064779 | -4.874387 | 1.09E-06 | 0.000232 |
| Sbf1 | 3831.593 | -0.315656 | 0.067031 | -4.70911 | 2.49E-06 | 0.000438 |
| Tmem145 | 2385.42 | -0.313594 | 0.078455 | -3.997115 | 6.41E-05 | 0.004591 |
| Luc7l2 | 9801.547 | -0.313066 | 0.067114 | -4.664716 | 3.09E-06 | 0.000523 |
| Unc13b | 1554.469 | -0.30997 | 0.076662 | -4.043356 | 5.27E-05 | 0.004055 |
| Plekha5 | 2605.094 | -0.307603 | 0.068229 | -4.508415 | 6.53E-06 | 0.000902 |
| Clk4 | 2531.849 | -0.30664 | 0.075018 | -4.087562 | 4.36E-05 | 0.003538 |
| Prpf38b | 3352.555 | -0.306141 | 0.058911 | -5.196678 | 2.03E-07 | 6.47E-05 |
| Zfp445 | 7601.274 | -0.305009 | 0.066305 | -4.60007 | 4.22E-06 | 0.000658 |
| Arhgef25 | 3525.098 | -0.304441 | 0.057642 | -5.281571 | 1.28E-07 | 4.40E-05 |
| Ipo9 | 5619.438 | -0.302395 | 0.055922 | -5.407452 | 6.39E-08 | 2.59E-05 |
| Zdhhc8 | 2189.691 | -0.299991 | 0.062834 | -4.774346 | 1.80E-06 | 0.00036 |
| Zfyve28 | 1402.218 | -0.298566 | 0.072323 | -4.128216 | 3.66E-05 | 0.003167 |
| Fnbp4 | 2895.692 | -0.294511 | 0.070629 | -4.169823 | 3.05E-05 | 0.002815 |
| Sfswap | 2859.407 | -0.293188 | 0.061323 | -4.781045 | 1.74E-06 | 0.000359 |
| A230050P1 | 2485.109 | -0.291871 | 0.046743 | -6.244107 | 4.26E-10 | 5.19E-07 |
| Kcnab3 | 2758.997 | -0.291756 | 0.073327 | -3.978849 | 6.92E-05 | 0.004755 |
| Pfkfb2 | 1363.106 | -0.279017 | 0.065067 | -4.288129 | 1.80E-05 | 0.001885 |
| Gigyf1 | 2930.245 | -0.278831 | 0.05867 | -4.752564 | 2.01E-06 | 0.000379 |
| Nrbp2 | 4941.377 | -0.278821 | 0.058759 | -4.745199 | 2.08E-06 | 0.000387 |
| Dpysl4 | 1383.457 | -0.270159 | 0.062596 | -4.315945 | 1.59E-05 | 0.00175 |
| Ubp1 | 3748.36 | -0.269475 | 0.059126 | -4.557629 | 5.17E-06 | 0.000753 |
| Dus3l | 2049.174 | -0.265019 | 0.061021 | -4.343095 | 1.40E-05 | 0.001622 |
| Ddx17 | 14279.92 | -0.263966 | 0.05065 | -5.211518 | 1.87E-07 | 6.12E-05 |
| Fbf1 | 2445.094 | -0.262705 | 0.057779 | -4.546706 | 5.45E-06 | 0.000776 |
| Lmtk3 | 6239.08 | -0.262413 | 0.044285 | -5.925498 | 3.11E-09 | 2.61E-06 |
| Kif21b | 3413.82 | -0.262336 | 0.058614 | -4.475662 | 7.62E-06 | 0.00102 |
| Gnl2 | 2292.289 | -0.258176 | 0.060751 | -4.249725 | 2.14E-05 | 0.002188 |
| **Celsr2** | 6982.925 | -0.257933 | 0.048364 | -5.33318 | 9.65E-08 | 3.49E-05 |
| Shc2 | 1404.425 | -0.254419 | 0.058788 | -4.327723 | 1.51E-05 | 0.001695 |
| Atxn2l | 5550.269 | -0.253305 | 0.058658 | -4.318332 | 1.57E-05 | 0.00175 |
| Cabin1 | 2452.666 | -0.2498 | 0.062298 | -4.009764 | 6.08E-05 | 0.004496 |
| Tecpr1 | 3272.945 | -0.248537 | 0.054312 | -4.57607 | 4.74E-06 | 0.000723 |
| Cables2 | 1004.409 | -0.24834 | 0.062371 | -3.981633 | 6.84E-05 | 0.004748 |
| **Irak1** | 1815.299 | -0.248294 | 0.054943 | -4.519085 | 6.21E-06 | 0.000866 |
| Agrn | 4411.07 | -0.247441 | 0.048171 | -5.136766 | 2.80E-07 | 8.04E-05 |
| Grk6 | 1346.402 | -0.245101 | 0.052648 | -4.655478 | 3.23E-06 | 0.000534 |
| Srrt | 3667.368 | -0.244421 | 0.054673 | -4.470569 | 7.80E-06 | 0.001034 |
| Haghl | 1862.053 | -0.242702 | 0.055538 | -4.369998 | 1.24E-05 | 0.001459 |
| Pabpn1 | 3658.565 | -0.242005 | 0.049775 | -4.861987 | 1.16E-06 | 0.000243 |
| Dgkz | 11126.44 | -0.238754 | 0.054564 | -4.375654 | 1.21E-05 | 0.001435 |
| Bbs7 | 1208.803 | -0.234117 | 0.053884 | -4.344788 | 1.39E-05 | 0.001622 |
| Ralgps1 | 2635.12 | -0.228667 | 0.056875 | -4.020545 | 5.81E-05 | 0.004343 |
| Ewsr1 | 7703.614 | -0.22525 | 0.054529 | -4.13085 | 3.61E-05 | 0.003163 |
| Sun1 | 3055.376 | -0.223208 | 0.049356 | -4.52242 | 6.11E-06 | 0.000862 |
| Srcin1 | 10952.21 | -0.211091 | 0.051392 | -4.107438 | 4.00E-05 | 0.003383 |
| Fbxl20 | 2732.773 | -0.209457 | 0.046937 | -4.462523 | 8.10E-06 | 0.001063 |
| B4galnt4 | 2330.951 | -0.209044 | 0.052171 | -4.006915 | 6.15E-05 | 0.004501 |
| Tnrc6a | 5065.66 | -0.20721 | 0.050008 | -4.143558 | 3.42E-05 | 0.003053 |
| Snx32 | 1837.339 | -0.202572 | 0.0478 | -4.237926 | 2.26E-05 | 0.002254 |
| Gripap1 | 3762.123 | -0.201953 | 0.047806 | -4.224419 | 2.40E-05 | 0.002356 |
| Ptprs | 10842.67 | -0.197215 | 0.043248 | -4.560158 | 5.11E-06 | 0.000752 |
| Kifc2 | 13045.44 | -0.190228 | 0.034956 | -5.44185 | 5.27E-08 | 2.21E-05 |
| Caskin1 | 4707.147 | -0.188713 | 0.042076 | -4.485092 | 7.29E-06 | 0.000986 |
| Rab1a | 6036.962 | 0.141288 | 0.03228 | 4.377015 | 1.20E-05 | 0.001435 |
| Hsp90ab1 | 44501.34 | 0.158378 | 0.037602 | 4.212019 | 2.53E-05 | 0.002404 |
| Arl6ip1 | 6076.481 | 0.160555 | 0.039481 | 4.066607 | 4.77E-05 | 0.003757 |
| Tmem184c | 3110.079 | 0.17744 | 0.044034 | 4.029632 | 5.59E-05 | 0.004202 |
| Saraf | 7738.089 | 0.17977 | 0.044394 | 4.049386 | 5.14E-05 | 0.003998 |
| Nus1 | 3156.129 | 0.194786 | 0.046313 | 4.205848 | 2.60E-05 | 0.002452 |
| Rps15a | 3590.375 | 0.199204 | 0.050176 | 3.970084 | 7.18E-05 | 0.004883 |
| Smarca2 | 11829.12 | 0.202383 | 0.050853 | 3.979762 | 6.90E-05 | 0.004755 |
| Ptges3 | 3869.774 | 0.206728 | 0.042232 | 4.895038 | 9.83E-07 | 0.000216 |
| Mapk4 | 2256.797 | 0.211486 | 0.05284 | 4.002406 | 6.27E-05 | 0.004538 |
| Itgb8 | 2611.055 | 0.212846 | 0.05221 | 4.076741 | 4.57E-05 | 0.003662 |
| Stip1 | 5720.957 | 0.213171 | 0.053378 | 3.993602 | 6.51E-05 | 0.004629 |
| Atp1a2 | 27999.43 | 0.217491 | 0.053741 | 4.047031 | 5.19E-05 | 0.004015 |
| Abhd3 | 3070.321 | 0.217773 | 0.044529 | 4.890559 | 1.01E-06 | 0.000217 |
| Elovl5 | 3578.33 | 0.224615 | 0.050911 | 4.411932 | 1.02E-05 | 0.001294 |
| Cnih1 | 1456.796 | 0.229083 | 0.046745 | 4.900731 | 9.55E-07 | 0.000215 |
| Dnajc3 | 2609.384 | 0.231514 | 0.056162 | 4.122269 | 3.75E-05 | 0.0032 |
| Acot11 | 2079.041 | 0.244257 | 0.051154 | 4.774956 | 1.80E-06 | 0.00036 |
| Paqr8 | 5979.539 | 0.246327 | 0.055537 | 4.435358 | 9.19E-06 | 0.001191 |
| Gm20594 | 17670.03 | 0.247728 | 0.060389 | 4.102217 | 4.09E-05 | 0.003405 |
| Kcnj10 | 12883.76 | 0.253446 | 0.051069 | 4.962827 | 6.95E-07 | 0.000164 |
| Prex2 | 2512.885 | 0.254995 | 0.059978 | 4.251469 | 2.12E-05 | 0.002187 |
| Tmem64 | 1400.639 | 0.256619 | 0.063671 | 4.030404 | 5.57E-05 | 0.004202 |
| Pdia3 | 6756.918 | 0.257778 | 0.064234 | 4.013126 | 5.99E-05 | 0.004457 |
| Chst2 | 3223.714 | 0.262252 | 0.060446 | 4.338594 | 1.43E-05 | 0.001641 |
| Ncoa4 | 1527.523 | 0.263299 | 0.064955 | 4.053548 | 5.04E-05 | 0.00395 |
| Rab8b | 1830.372 | 0.270176 | 0.067044 | 4.029843 | 5.58E-05 | 0.004202 |
| Gja1 | 8138.44 | 0.274291 | 0.054053 | 5.074496 | 3.89E-07 | 9.82E-05 |
| Hnrnpc | 7793.682 | 0.281739 | 0.064108 | 4.394739 | 1.11E-05 | 0.00135 |
| Anxa5 | 3975.859 | 0.28249 | 0.066102 | 4.27357 | 1.92E-05 | 0.001997 |
| mt-Atp8 | 57608.54 | 0.298342 | 0.04715 | 6.327502 | 2.49E-10 | 4.17E-07 |
| Hsp90b1 | 14215.27 | 0.298935 | 0.058423 | 5.116749 | 3.11E-07 | 8.33E-05 |
| Insig2 | 1193.582 | 0.310298 | 0.075803 | 4.09348 | 4.25E-05 | 0.003491 |
| Lrp4 | 980.569 | 0.325328 | 0.075712 | 4.296908 | 1.73E-05 | 0.001826 |
| Gabpa | 1332.866 | 0.328273 | 0.080618 | 4.071982 | 4.66E-05 | 0.003693 |
| Tmem47 | 4623.177 | 0.339744 | 0.082975 | 4.094536 | 4.23E-05 | 0.003491 |
| Rgs5 | 2606.863 | 0.341233 | 0.072547 | 4.703604 | 2.56E-06 | 0.000441 |
| Stt3a | 1609.708 | 0.348354 | 0.056204 | 6.197994 | 5.72E-10 | 6.38E-07 |
| Bag3 | 546.2413 | 0.362319 | 0.071942 | 5.036238 | 4.75E-07 | 0.000118 |
| Rab9b | 1431.103 | 0.373848 | 0.08848 | 4.225206 | 2.39E-05 | 0.002356 |
| Ddx3x | 15060.22 | 0.388105 | 0.076048 | 5.103445 | 3.34E-07 | 8.76E-05 |
| Vamp7 | 1125.509 | 0.397276 | 0.060768 | 6.53758 | 6.25E-11 | 1.40E-07 |
| Slc7a11 | 1740.342 | 0.398035 | 0.089914 | 4.426819 | 9.56E-06 | 0.00122 |
| Calr | 14324.86 | 0.402171 | 0.088149 | 4.562393 | 5.06E-06 | 0.000752 |
| **Kcnj16** | 1618.676 | 0.406042 | 0.08526 | 4.762368 | 1.91E-06 | 0.000371 |
| Pcdhb14 | 580.0079 | 0.40846 | 0.072492 | 5.634555 | 1.76E-08 | 8.39E-06 |
| Tsc22d3 | 1421.047 | 0.408729 | 0.089602 | 4.561614 | 5.08E-06 | 0.000752 |
| Fam107a | 9772.404 | 0.414724 | 0.096107 | 4.315223 | 1.59E-05 | 0.00175 |
| Hspa5 | 8390.226 | 0.43407 | 0.093739 | 4.630625 | 3.65E-06 | 0.000588 |
| Rps12 | 952.2666 | 0.463544 | 0.110984 | 4.176669 | 2.96E-05 | 0.002751 |
| Efemp1 | 546.3291 | 0.482194 | 0.09719 | 4.961337 | 7.00E-07 | 0.000164 |
| mt-Nd6 | 3292.188 | 0.514203 | 0.111346 | 4.618053 | 3.87E-06 | 0.000617 |
| Aebp1 | 577.8293 | 0.524505 | 0.12364 | 4.242191 | 2.21E-05 | 0.002228 |
| Wnt5a | 396.5741 | 0.527424 | 0.119903 | 4.398738 | 1.09E-05 | 0.001338 |
| Colec12 | 601.0315 | 0.551464 | 0.108406 | 5.087032 | 3.64E-07 | 9.37E-05 |
| Eif2s3x | 5856.155 | 0.558882 | 0.095169 | 5.872516 | 4.29E-09 | 3.02E-06 |
| Mt2 | 1590.433 | 0.56403 | 0.089852 | 6.277319 | 3.44E-10 | 4.61E-07 |
| Nrros | 151.3606 | 0.580683 | 0.141905 | 4.092046 | 4.28E-05 | 0.003491 |
| Dcn | 505.3374 | 0.604025 | 0.147831 | 4.085915 | 4.39E-05 | 0.003541 |
| Tnfrsfm13 | 168.0315 | 0.615449 | 0.14916 | 4.126091 | 3.69E-05 | 0.003167 |
| Ctsh | 284.4134 | 0.627951 | 0.115173 | 5.45222 | 4.97E-08 | 2.15E-05 |
| Ccdc36 | 255.7494 | 0.705423 | 0.153029 | 4.609731 | 4.03E-06 | 0.000635 |
| Spp1 | 529.2915 | 0.719683 | 0.162308 | 4.434071 | 9.25E-06 | 0.001191 |
| Cyp1b1 | 184.0243 | 0.72518 | 0.174787 | 4.148921 | 3.34E-05 | 0.003033 |
| Col1a2 | 327.8589 | 0.737056 | 0.143895 | 5.122191 | 3.02E-07 | 8.33E-05 |
| **Avp** | 323.053 | 0.783328 | 0.192355 | 4.072303 | 4.66E-05 | 0.003693 |
| Ogn | 232.7831 | 0.875971 | 0.185755 | 4.71573 | 2.41E-06 | 0.00043 |
| Emp1 | 125.7175 | 0.894582 | 0.18259 | 4.8994 | 9.61E-07 | 0.000215 |

Note: Genes in bold text appear in the DEGs between AAV-GFP and WT mice.
Also note: Only DEGs with Padj values < 0.05 are listed; down-regulated and up-regulated genes in AAV-lncRNA mutants in blue and red text respectively.
Low and high dose lncRNA-treated mice form a single group for the analysis.

**Supplemental Table 4.**
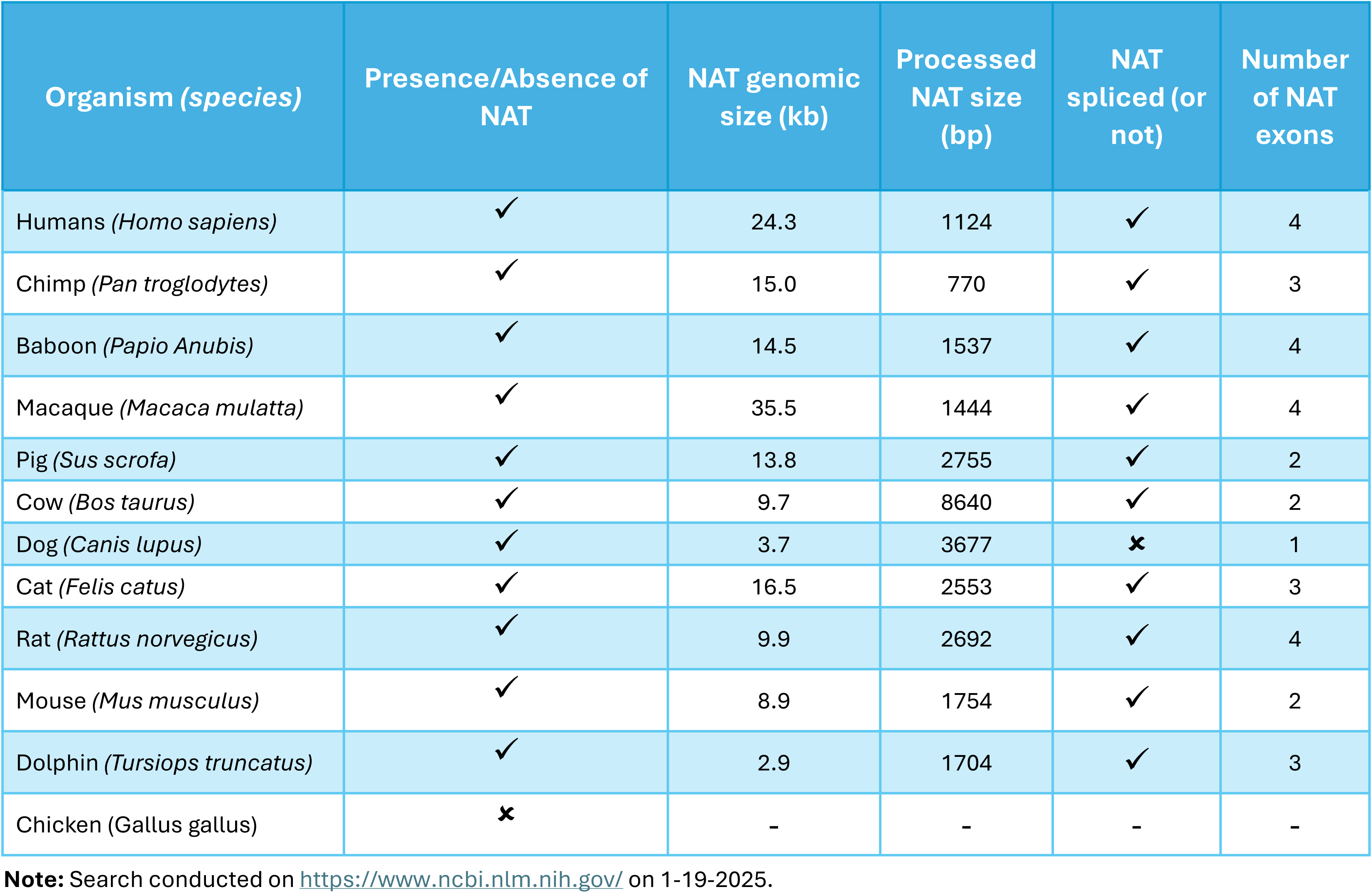
Prevalence and characteristics of natural antisense transcripts in the *SLC2A1 (Glut1)* locus of diverse species.

